## Supplemental for "Conformational changes in the essential *E. coli* septal cell wall synthesis complex suggest an activation mechanism"

**Supplementary Table S1:** Plasmids used in this study. Descriptions of the plasmid components and overviews of construction methods are shown below.

| PLASMID | GENOTYPE | SOURCE |
|---|---|---|
| **PBMB040** | pCH-HaloTag-FtsB (Kan) | This study |
| **PBMB053** | pCH-HaloTag-FtsB (Carb) | This study |
| **PJM090** | FtsI WT (Kan) | This study |
| **PJM092** | FtsI Δ10 (Kan) | This study |
| **PJM093** | FtsI Δ14 (Kan) | This study |
| **PBMB063** | FtsL WT (Carb) | This study |
| **PBMB064** | FtsL Δ6 (Carb) | This study |
| **PBMB065** | FtsL Δ11 (Carb) | This study |
| **PBMB066** | FtsL Δ16 (Carb) | This study |
| **PBMB068** | pVS155-FtsL | [1] |
| **PBMB069** | Venus-FtsL WT (Carb) | This study |
| **PBMB070** | Venus-FtsL Δ6 (Carb) | This study |
| **PBMB071** | Venus-FtsL Δ11 (Carb) | This study |
| **PBMB072** | Venus-FtsL Δ16 (Carb) | This study |

**pBMB040:** The *ftsB* gene was amplified from *E. coli* genomic DNA using primer pair oBMB027 and oBMB076. The pCH-Kan vector backbone containing the *HaloTag* gene was amplified using primer pair oBMB077 and oBMB078. The two DNA fragments were combined using In-Fusion cloning to generate pBMB040 (HaloTag-FtsB (Kan)).

**pBMB053:** The *HaloTag-ftsB* gene was amplified from pBMB040 using oBMB077 and oBMB016. The vector backbone pCH-Carb was amplified using oBMB010 and oBMB009. The two DNA fragments were combined using In-Fusion cloning to generate pBMB053 (HaloTag-FtsB (Carb)).

**pJM090:** The *ftsI* gene was amplified from *E. coli* genomic DNA with oJM288 and oJM229. The vector backbone pCH-Kan was amplified using oRM100 and oJM6. The two DNA fragments were combined using In-Fusion cloning generating pJM090 (FtsI WT (Kan)).

**pJM092:** The plasmid pLM090 was reverse amplified using oJM230 and oJM231 and circularized using In-Fusion cloning to generate pJM092 that contains the truncated *ftsI* gene (FtsI Δ10 (Kan)).

**pJM093:** The plasmid pLM090 was reverse amplified using oJM234 and oJM235 and circularized using In-Fusion cloning to generate pJM093 that contains the truncated *ftsI* gene (FtsI Δ14 (Kan)).

**pBMB063:** The *ftsL* gene was amplified from *E. coli* genomic DNA using oBMB013 and oBMB014. The vector backbone pCH-Carb was amplified using oBMB010 and oBMB009. The two DNA fragments were combined using In-Fusion cloning generating pBMB063 (FtsL WT (Carb)).

**pBMB064:** The plasmid pBMB063 was reverse-amplified using oBMB121 and oBMB122 and circularized using In-Fusion cloning to generate pBMB064 that contains a truncated *ftsL* gene (FtsL Δ6 (Carb)).

**pBMB065:** The plasmid pBMB063 was reverse amplified using oBMB121 and oBMB123 and circularized using In-Fusion cloning to generate pBMB065 that contains a truncated *ftsL* gene (FtsL Δ11 (Carb)).

**pBMB066:** The plasmid pBMB063 was reverse amplified using oBMB121 and oBMB124 and circularized using In-Fusion cloning to generate pBMB066 that contains a truncated *ftsL* gene (FtsL Δ16 (Carb)).

**pBMB068:** The plasmid pBMB068 was the same as pVS155-FtsL in ref [1] but renamed to maintain the naming consistency of the current work.

**pBMB069:** The *mVenus* gene was amplified from pBMB068 using oBMB127 and oBMB128. The *ftsL* gene and vector backbone were amplified from pBMB063 using oBMB010 and oBMB009. The two DNA fragments were combined using In-Fusion cloning to generate pBMB069 (Venus-FtsL WT (Carb)).

**pBMB070:** The *mVenus* gene was amplified from pBMB068 using oBMB127 and oBMB128. The truncated *ftsL* gene and vector backbone were amplified from pBMB064 using oBMB010 and oBMB122. The two DNA fragments were combined using In-Fusion cloning to generate pBMB069 (Venus-FtsL Δ6 (Carb)).

**pBMB071:** The *mVenus* gene was amplified from pBMB068 using oBMB127 and oBMB128. The truncated *ftsL* gene and vector backbone were amplified from pBMB065 using oBMB010 and oBMB123. The two DNA fragments were combined using In-Fusion cloning to generate pBMB071 (Venus-FtsL Δ11 (Carb)).

**pBMB072:** The *mVenus* gene was amplified from pBMB068 using oBMB127 and oBMB128. The truncated *ftsL* gene and vector backbone were amplified from pBMB066 using oBMB010 and oBMB124. The two DNA fragments were combined using In-Fusion cloning to generate pBMB072 (Venus-FtsL Δ72 (Carb)).

**Supplementary Table S2:** Strains used in this study.

| Strains | Genotype | Source |
|---|---|---|
| **TB28** | MG1655 *laclZYA < >frt* | [2] |
| **TM6** | TB28 *ftsIR167S* | [3] |
| **MDG279** | JOE309 Δ*ftsL::kan*/ *pBAD33-ftsL* | [4] |
| **CR14** | JOE309 *ΔftsB(Kanr)/pBAD42-ftsB* | [5] |
| **EC812** | MG1655 *ftsI::cat Δ(λattL-lom)::bla lacIq pBAD-ftsI* | [6] |
| **sBMB015** | TB28 / *pCH-halo-ftsB* (Kan) | This study. Transform pBMB040 into TB28, select Kan$^{r}$. |
| **sBMB052** | TM6 / *pCH-halo-ftsB* (Kan) | This study. Transform pBMB040 into TM6, select Kan$^{r}$. |
| **sBMB184** | CR14 / *pCH-halo-ftsB* (Carb) | This study. Transform pBMB053 into CR14, select Carb$^{r}$. |
| **sBMB121** | EC812 / *pCH-ftsI WT* (Kan) | This study. Transform pJM90 into EC812, select Kan$^{r}$. |
| **sBMB122** | EC812 / *ftsI Δ10* (Kan) | This study. Transform pJM092 into EC812, select Kan$^{r}$. |
| **sBMB123** | EC812 / *ftsI Δ14* (Kan) | This study. Transform pJM093 into EC812, select Kan$^{r}$. |
| **sBMB097** | MDG279 / *ftsL Δ11* (Carb) | This study. Transform pBMB065 into MDG279, select Carb$^{r}$. |
| **sBMB099** | MDG279 / *ftsL Δ16* (Carb) | This study. Transform pBMB066 into MDG279, select Carb$^{r}$. |
| **sBMB103** | MDG279 / *ftsL Δ6* (Carb) | This study. Transform pBMB064 into MDG279, select Carb$^{r}$. |
| **sBMB124** | MDG279 / *ftsL wt* (Carb) | This study. Transform pBMB063 into MDG279, select Carb$^{r}$. |
| **sBMB146** | MDG279 / *venus -ftsL wt* (Carb) | This study. Transform pBMB069 into MDG279, select Carb$^{r}$. |
| **sBMB147** | MDG279 / *venus -ftsL Δ6* (Carb) | This study. Transform pBMB070 into MDG279, select Carb$^{r}$. |
| **sBMB148** | MDG279 / *venus -ftsL Δ11* (Carb) | This study. Transform pBMB071 into MDG279, select Carb$^{r}$. |
| **sBMB149** | MDG279 / *venus -ftsL Δ16* (Carb) | This study. Transform pBMB072 into MDG279, select Carb$^{r}$. |

**Supplementary Table S3:** FtsB dynamics under different sPG synthesis conditions.

| Genotype | Drug or medium | $P_{moving}$ (%), μ ± s.e.m., ($N_{all}$) | $P_{slow}$ (%), μ ± s.e.m. | $V_{slow}$ (nm/s), μ ± s.e.m., (N) | $V_{fast}$ (nm/s), μ ± s.e.m. |
|---|---|---|---|---|---|
| **TB28** | M9(-) glucose | 53.7% ± 2.7% (379) | 62% ± 13% | 8.2 ± 1.7 (179) | 32.9 ± 5.2 |
| **TB28** | Fosfomycin, M9(-) glucose | 44.1% ± 3.8% (495) | 30% ± 10% | 8.5 ± 3.0 (218) | 28.5 ± 2.9 |
| **TB28, *ftsI*$^{R167S}$** | EZRDM | 37.0% ± 3.1% (322) | 100% | 8.0 ± 0.4 (112) | N.A. |

**Supplementary Table S4.** Values of pDockQ were calculated using a script in PyMOL according to previously reported methods [7].The quality of the predicted protein-protein interfaces for each subunit to all others was estimated in this way.

| | FtsQLBWI | FtsQLBWIN | FtsQLBWIN$^{58–108}$ |
|---|---|---|---|
| **FtsQ** | 0.64 | 0.62 | 0.64 |
| **FtsL** | 0.73 | 0.73 | 0.73 |
| **FtsB** | 0.73 | 0.73 | 0.73 |
| **FtsW** | 0.72 | 0.72 | 0.72 |
| **FtsI** | 0.72 | 0.73 | 0.73 |
| **FtsN** | N/A | 0.61 | 0.61 |

**Supplementary Table S5.** Details of MD simulation systems.

| System | Dimensions | No. atoms | Duration |
|---|---|---|---|
| **FtsQLBWI (WT)** | 180 Å x 180 Å x 206 Å | 645896 | 1 μs |
| **FtsQLBWIN (FtsN)** | 180 Å x 180 Å x 209 Å | 653520 | 1 μs |
| **FtsQLBWI$^{R167S}$ (R167S)** | 180 Å x 180 Å x 209 Å | 654206 | 1 μs |
| **FtsQLBW$^{E289G}$I (E289G)** | 180 Å x 180 Å x 209 Å | 654426 | 1 μs |
| **FtsQL$^{R61E}$BWI (R61E)** | 180 Å x 180 Å x 206 Å | 645657 | 1 μs |
| **FtsWI** | 160 Å x 160 Å x 211 Å | 523966 | 1 μs |
| **FtsQLBWI (with truncated FtsI C-terminus)** | 180 Å x 180 Å x 196 Å | 614294 | 200 ns |

**Supplementary Table S6:** FtsL cell length distributions. All samples are strain MDG279 harboring plasmids expressing various FtsL constructs in different induction conditions.

| Strain and growth condition | Mean Length (μm, μ ± s.e.m.) | Number of Cells |
|---|---|---|
| ***ΔftsL*** | 13.1 ± 2.7 | 243 |
| ***ΔftsL* + *ftsL*** | 2.7 ± 0.1 | 1606 |
| ***ftsL wt* + 0 μM IPTG** | 5.6 ± 0.4 | 599 |
| ***ftsL wt* + 1 μM IPTG** | 4.5 ± 0.3 | 538 |
| ***ftsL wt* + 10 μM IPTG** | 3.8 ± 0.8 | 1027 |
| ***ftsL wt* + 100 μM IPTG** | 3.3 ± 0.2 | 436 |
| ***ftsL Δ6* + 0 μM IPTG** | 8.4 ± 1.6 | 121 |
| ***ftsL Δ6* + 1 μM IPTG** | 6.0 ± 1.1 | 443 |
| ***ftsL Δ6* + 10 μM IPTG** | 5.4 ± 0.9 | 389 |
| ***ftsL Δ6* + 100 μM IPTG** | 4.7 ± 0.7 | 417 |
| ***ftsL Δ11* + 0 μM IPTG** | 11.4 ± 4.8 | 213 |
| ***ftsL Δ11* + 1 μM IPTG** | 7.1 ± 2.1 | 248 |
| ***ftsL Δ11* + 10 μM IPTG** | 4.8 ± 1.5 | 163 |
| ***ftsL Δ11* + 100 μM IPTG** | 4.8 ± 0.7 | 724 |
| ***ftsL Δ16* + 0 μM IPTG** | 17.2 ± 10.2 | 86 |
| ***ftsL Δ16* + 1 μM IPTG** | 8.9 ± 3.4 | 113 |
| ***ftsL Δ16* + 10 μM IPTG** | 5.8 ± 0.4 | 117 |
| ***ftsL Δ16* + 100 μM IPTG** | 4.5 ± 0.6 | 278 |

**Supplementary Table S7:** FtsL Midcell Fluorescence Ratio. Fluorescence image quantification of the fraction of fluorescence intensity at midcell.

| Strain and growth condition | Percentage (μ ± s.d.) | Number of Cells |
|---|---|---|
| ***ftsL wt* + 100 μM IPTG** | 0.40 ± 0.08 | 225 |
| ***ftsL Δ6* + 100 μM IPTG** | 0.37 ± 0.08 | 178 |
| ***ftsL Δ11* + 100 μM IPTG** | 0.35 ± 0.08 | 177 |
| ***ftsL Δ16* + 100 μM IPTG** | 0.31 ± 0.07 | 144 |

**Supplementary Table S8:** FtsL Fluorescence/Cell Length Intensity. Fluorescence image quantification of total cellular fluorescence level normalized against cell length.

| Strain and growth condition | Intensity (a.u., μ ± s.d.) | Number of Cells |
|---|---|---|
| ***ftsL wt* + 100 μM IPTG** | 67 ± 21 | 225 |
| ***ftsL Δ6* + 100 μM IPTG** | 70 ± 25 | 178 |
| ***ftsL Δ11* + 100 μM IPTG** | 88 ± 25 | 177 |
| ***ftsL Δ16* + 100 μM IPTG** | 95 ± 25 | 144 |

**Supplementary Table S9:** FtsI cell length distributions. All samples are strain EC812 harboring various FtsI expression plasmids in different induction conditions.

| Strain and growth condition | Mean Length (μm, μ ± s.e.m.,) | Number of Cells |
|---|---|---|
| ***ΔftsI*** | 18.1 ± 4.1 | 192 |
| ***ΔftsI* + *ftsI*** | 2.5 ± 0.1 | 3582 |
| ***ftsI wt* + 0 μM IPTG** | 4.3 ± 0.3 | 524 |
| ***ftsI wt* + 1 μM IPTG** | 3.9 ± 0.0 | 1215 |
| ***ftsI wt* + 10 μM IPTG** | 3.5 ± 0.3 | 1233 |
| ***ftsI wt* + 100 μM IPTG** | 2.9 ± 0.0 | 1165 |
| ***ftsI Δ10* + 0 μM IPTG** | 7.2 ± 0.2 | 1252 |
| ***ftsI Δ10* + 1 μM IPTG** | 5.5 ± 1.3 | 446 |
| ***ftsI Δ10* + 10 μM IPTG** | 3.2 ± 0.1 | 1689 |
| ***ftsI Δ10* + 100 μM IPTG** | 3.0 ± 0.1 | 938 |
| ***ftsI Δ14* + 0 μM IPTG** | 3.5 ± 0.2 | 974 |
| ***ftsI Δ14* + 1 μM IPTG** | 3.3 ± 0.0 | 487 |
| ***ftsI Δ14* + 10 μM IPTG** | 3.2 ± 0.2 | 1181 |
| ***ftsI Δ14* + 100 μM IPTG** | 3.0 ± 0.3 | 1200 |

**Supplementary Table S10:** FtsI Western blot quantification showed that $FtsI^{\Delta 10}$ expression was lower than that for either FtsI or $FtsI^{\Delta 14}$ at equivalent induction levels, and that $FtsI^{\Delta 14}$ expression was marginally higher than that of FtsI. Except for the TB28 sample, samples are for EC812 harboring various FtsI expression plasmids. Measurements for individual samples in each replicate are normalized to intensity for condition *ΔftsI* + *ftsI*.

| Strain and growth condition | Relative Quantity (μ ± s.d,) | Replicates |
|---|---|---|
| **TB28** | 0.61 ± 0.01 | 3 |
| ***ΔftsI* + *ftsI*** | 1 | 3 |
| ***ΔftsI*** | 0.19± 0.04 | 3 |
| ***ftsI wt* + 0 μM IPTG** | 0.44 ± 0.2 | 3 |
| ***ftsI wt* + 1 μM IPTG** | 0.62 ± 0.2 | 3 |
| ***ftsI wt* + 10 μM IPTG** | 1.2 ± 0.1 | 3 |
| ***ftsI Δ10* + 0 μM IPTG** | 0.34± 0.2 | 3 |
| ***ftsI Δ10* + 1 μM IPTG** | 0.33 ± 0.1 | 3 |
| ***ftsI Δ10* + 10 μM IPTG** | 0.82 ± 0.2 | 3 |
| ***ftsI Δ14* + 0 μM IPTG** | 0.43 ± 0.3 | 3 |
| ***ftsI Δ14* + 1 μM IPTG** | 0.71 ± 0.2 | 3 |
| ***ftsI Δ14* + 10 μM IPTG** | 1.2 ± 0.2 | 3 |

**Supplementary Table S11:** FtsI midcell localization is quantified in anti-FtsI immunofluorescence images as the ratio of midcell to total cellular fluorescence.

| Strain | Ratio ($\mu \pm$ s.d,) | Number of Cells |
|---|---|---|
| ***TB28*** | 0.7 ± 0.1 | 244 |
| ***Δftsl* + *ftsI*** | 0.6 ± 0.1 | 220 |
| ***ftsI wt* + 0 μM IPTG** | 0.6 ± 0.1 | 236 |
| ***ftsI wt* + 1 μM IPTG** | 0.6 ± 0.1 | 208 |
| ***ftsI wt* + 10 μM IPTG** | 0.5 ± 0.1 | 224 |
| ***ftsI wt* + 100 μM IPTG** | 0.5 ± 0.1 | 220 |
| ***ftsI Δ10* + 0 μM IPTG** | 0.6± 0.1 | 209 |
| ***ftsI Δ10* + 1 μM IPTG** | 0.6 ± 0.1 | 209 |
| ***ftsI Δ10* + 10 μM IPTG** | 0.5 ± 0.1 | 204 |
| ***ftsI Δ10* + 100 μM IPTG** | 0.5 ± 0.1 | 205 |
| ***ftsI Δ14* + 0 μM IPTG** | 0.6 ± 0.1 | 210 |
| ***ftsI Δ14* + 1 μM IPTG** | 0.6 ± 0.1 | 201 |
| ***ftsI Δ14* + 10 μM IPTG** | 0.6 ± 0.1 | 225 |
| ***ftsI Δ14* + 100 μM IPTG** | 0.5 ± 0.1 | 209 |

**Supplementary Table S12:** Oligonucleotides used in this study.

| Oligo | Sequence 5'- 3' | Purpose |
|---|---|---|
| **oJM6** | GCGGCCGCTAAGGGTCG | Amplification of vector backbone. |
| **oBMB009** | GCGGCCGCTAAGGGT | Amplification of *ftsL*. |
| **oBMB010** | ACTAGTAGTTAATTTCTCCTCTTTAATGAATTCTGTGTG | Amplification of *ftsL*. |
| **oBMB013** | AAATTAACTACTAGTATGATCAGCAGAGTGACAGAAGCTCTAAGCAAAGT | Amplification of vector backbone. |
| **oBMB014** | ACCCTTAGCGGCCGCTTATTTTTGCACTACGATATTTTCTTGTGACGGATCAAC ATGC | Amplification of vector backbone. |
| **oBMB016** | ACCCTTAGCGGCCGCTTATCGATTGTTTTGCCCCGCAGACTGT | Amplification of vector backbone. |
| **oBMB027** | ACTAGTAGTTAATTTCTCCTCTTTAATGAATTCTGTGTG | Amplification of *ftsB*. |
| **oBMB076** | ATTAACAAATTTGTGCAGAAAAGCCGTC | Amplification of *ftsB*. |
| **oBMB077** | AAATTAACTACTAGTATGGGATCCGAAATCGG | Amplification of vector backbone. |
| **oBMB078** | CACAAATTTGTTAATGGAAATCTCCAGAGTAGACAGCCAG | Amplification of vector backbone. |
| **oRM100** | ACTAGTAGTTAATTTCTCCTCTTTAATG | Amplification of vector backbone. |
| **oBMB121** | TAAGCGGCCGCTAAGGGT | Amplification of *ftsL*. |
| **oBMB122** | CTTAGCGGCCGCTTATTCTTGTGACGGATCAACATGCTG | Amplification of *ftsL* truncation. |
| **oBMB123** | CTTAGCGGCCGCTTAAACATGCTGCATTTGCAGCTTTTC | Amplification of *ftsL* truncation. |
| **oBMB124** | CTTAGCGGCCGCTTACAGCTTTTCCGTGGCGATCC | Amplification of *ftsL* truncation. |
| **oBMB127** | AAATTAACTACTAGTATGGTGAGCAAGGGCGAGGAGC | Amplification of mVenus. |
| **oBMB128** | CCTTTAACTTTGCTTAGAGCTTCTGTCACTCTGCTGAT | Amplification of mVenus. |
| **oJM228** | ACCCTTAGCGGCCGCTTACGATCTGCCACCTGTCC | Amplification of *ftsI.* |
| **oJM229** | AAATTAACTACTAGTATGAAAGCAGCGGCGAAAACG | Amplification of *ftsI.* |
| **oJM230** | CTTAGCGGCCGCTTAAATCACAAATTCATTTTTATCGCCCG | Amplification of *ftsI truncation.* |
| **oJM231** | AATGAATTTGTGATTTAAGCGGCCGCTAAGGGTC | Amplification of *ftsI* truncation. |
| **oJM234** | ACGGGCGATAAAAATTAAGCGGCCGCTAAGGGTC | Amplification of *ftsI* truncation. |
| **oJM235** | CTTAGCGGCCGCTTAATTTTTATCGCCCGTTGTCAGCG | Amplification of *ftsI* truncation. |

### Supplementary Figure S1

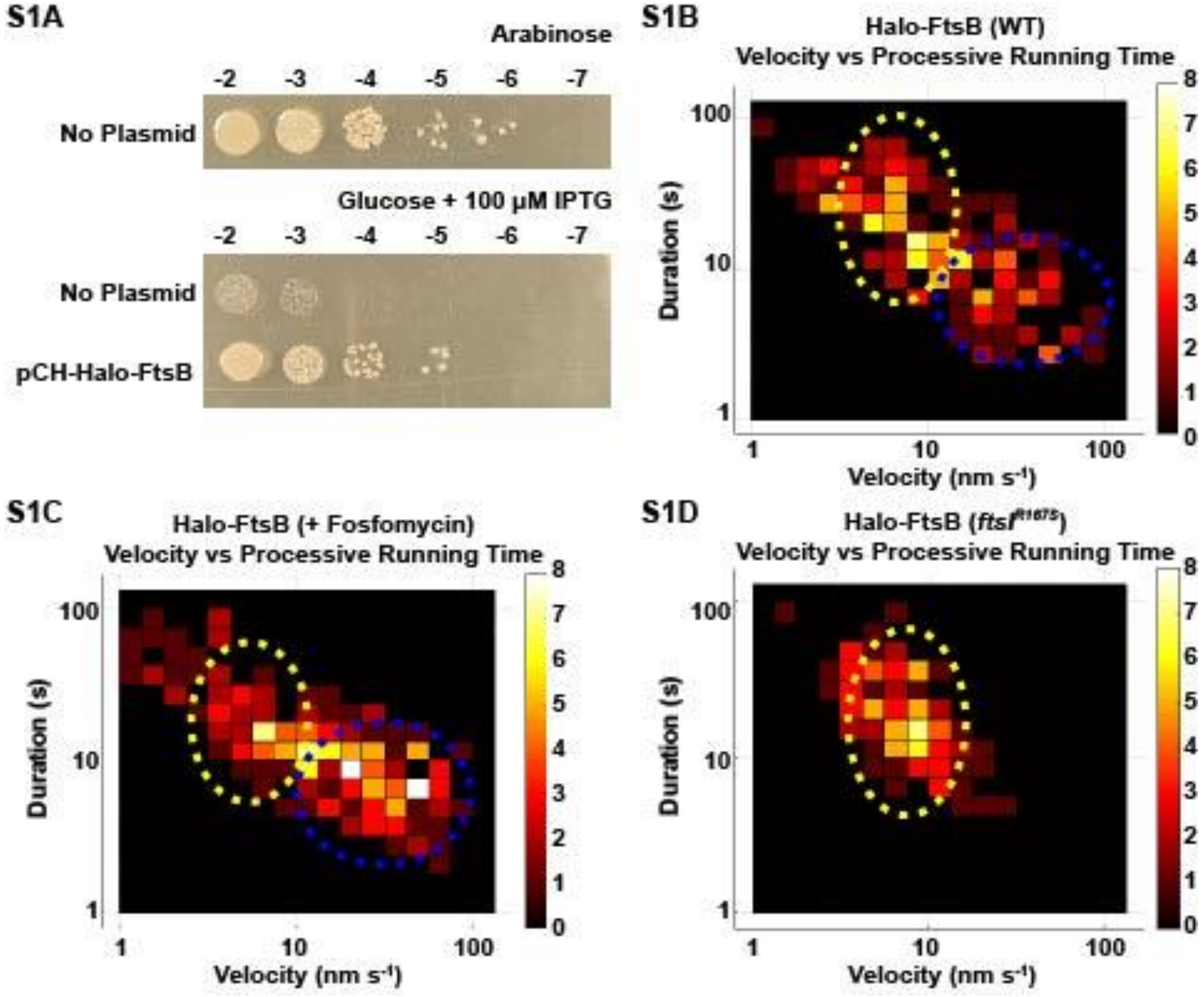


**Supplementary Figure S1.** **(A)** A Halo-FtsB fusion protein complemented an *E. coli* strain depleted of wild-type *ftsB*. FtsB depletion strain CR14 (**Table S2**) grew in the presence of arabinose, which induces FtsB expression from a $P_{BAD}$ promoter (pBAD-FtsB, top panel), but failed to form colonies in the absence of inducer (glucose + 100 μM IPTG panel, pBAD-FtsB, top row of the second panel). Expression of Halo-FtsB from plasmid pBMB053 (**Table S1**) complemented FtsB depletion in CR14 when induced with 100 μM IPTG (pCH-Halo-FtsB, bottom row of second panel). **(B to D)** Two-dimensional histograms of the directional moving velocity (Velocity, x axis) and processive running time (Duration, y axis) of single Halo-FtsB molecules expressed from plasmid pBMB040 in WT (**B**), in WT cells treated with fosfomycin (**C**), and in the *ftsI*$^{R167S}$ superfission mutant background (**D**). The color bars indicate the frequency of each bin. The two moving populations, one fast (blue dotted oval) and one slow (yellow dotted oval) were observed in WT (**B**). The two populations shifted to a higher fraction of the fast-moving population (blue dotted oval) when treated with fosfomycin (**C**). The fast-moving population nearly disappeared and the slow moving population (yellow dotted oval) dominated in the *ftsI*$^{R167S}$ superfission mutant background **(D**). These changes are similar to what we observed for FtsW [3].

### Supplementary Figure S2

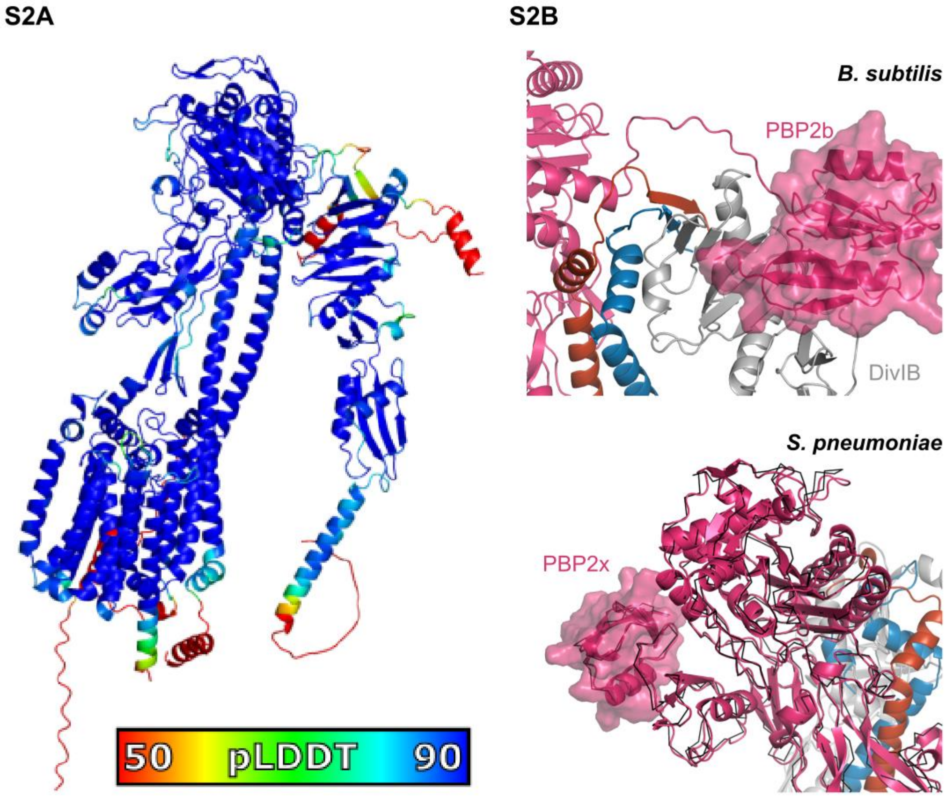


**Supplementary Figure S2. (A)** Predicted AF2 structure of full-length *E. coli* FtsQLBWI, colored by local prediction confidence, pLDDT. Red regions with pLDDT below 50 are $\text{FtsQ}^{1-24}$, $\text{FtsQ}^{1-24}$, $\text{FtsQ}^{263-276}$, $\text{FtsL}^{1-21}$, $\text{FtsB}^{92-103}$, $\text{FtsW}^{1-40}$, $\text{FtsW}^{413-414}$, $\text{FtsI}^{1-17}$, and $\text{FtsI}^{580-588}$. **(B)** Predictions of homologous FtsQLBWI complexes in *B. subtilis* and *S. pneumonia*, zooming in on regions with previously reported interactions for C-terminal PASTA domains for divisomal PBPs. Top, C-terminal PASTA domain of *B. subtilis* PBP2b (magenta) is predicted to bind DivIB (gray, FtsQ in *E. coli*) as observed in bacterial two-hybrid and co-immunoprecipitation assays [8]. Bottom, the C-terminal PASTA domain in *S. pneumoniae* PBP2x in the predicted complex adopts the same structure previously reported (black lines from PDB 5OIZ) to have allosteric autoregulatory activity [9].

### Supplementary Figure S3

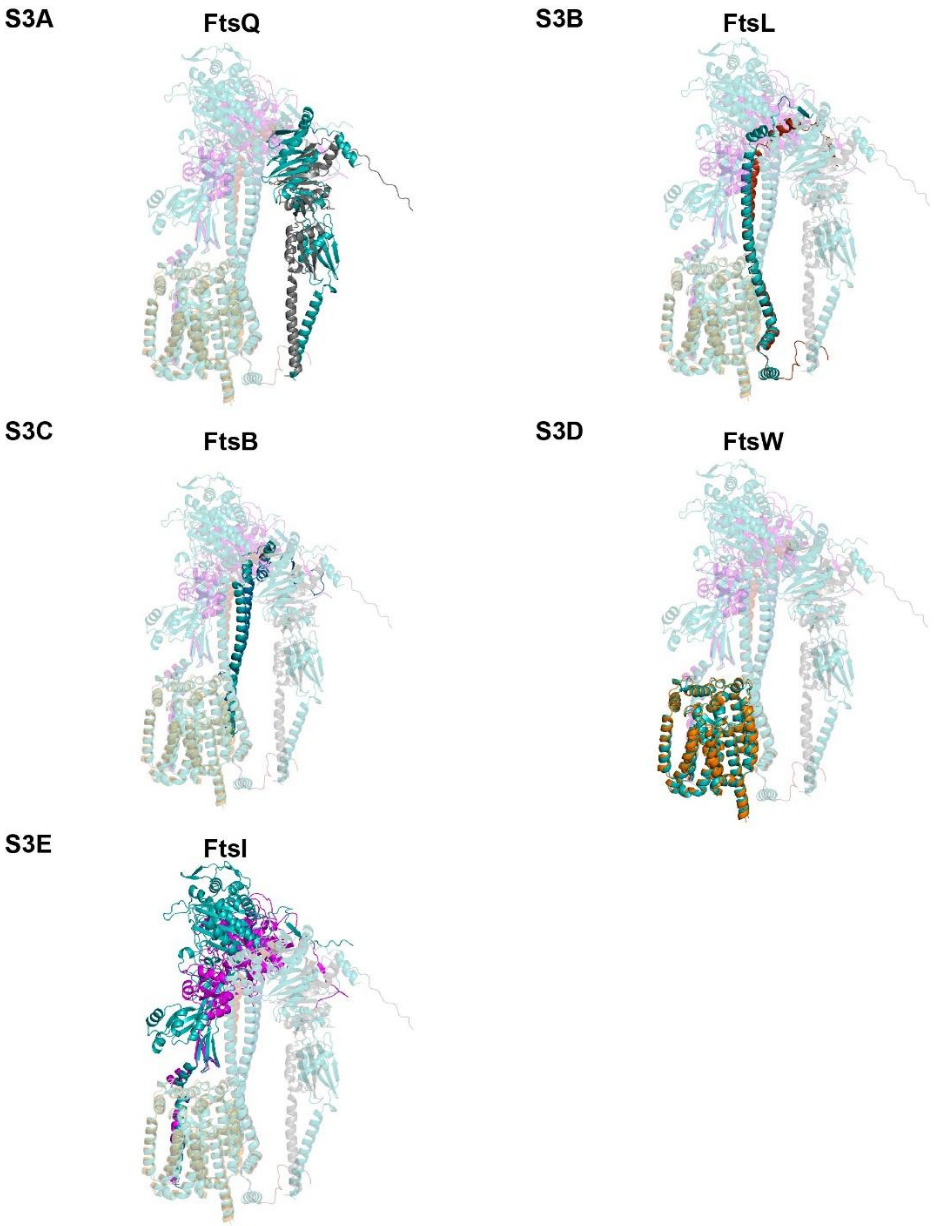


**Supplementary Figure S3.** Evolution of AF2 FtsQLBWI model before (teal) and after MD simulation (colored) for each protein**.** For each protein, the conformer after 1 µs of MD simulation is shown for **(A)** FtsQ (gray) **(B)** FtsL (red) **(C)** FtsB (blue) **(D)** FtsW (orange) **(E)** FtsI (magenta). Other proteins not in comparison in each complex are shown with transparent representations. The same orientation is shown in all subfigures with coordinates for predicted structures and after 1-µs MD aligned by FtsW. PDBs for predicted structures and coordinates after 1-µs MD are available for download. Major changes during MD were observed for the orientation of the FtsI TPase domain relative to FtsW (**E**).

### Supplementary Figure S4

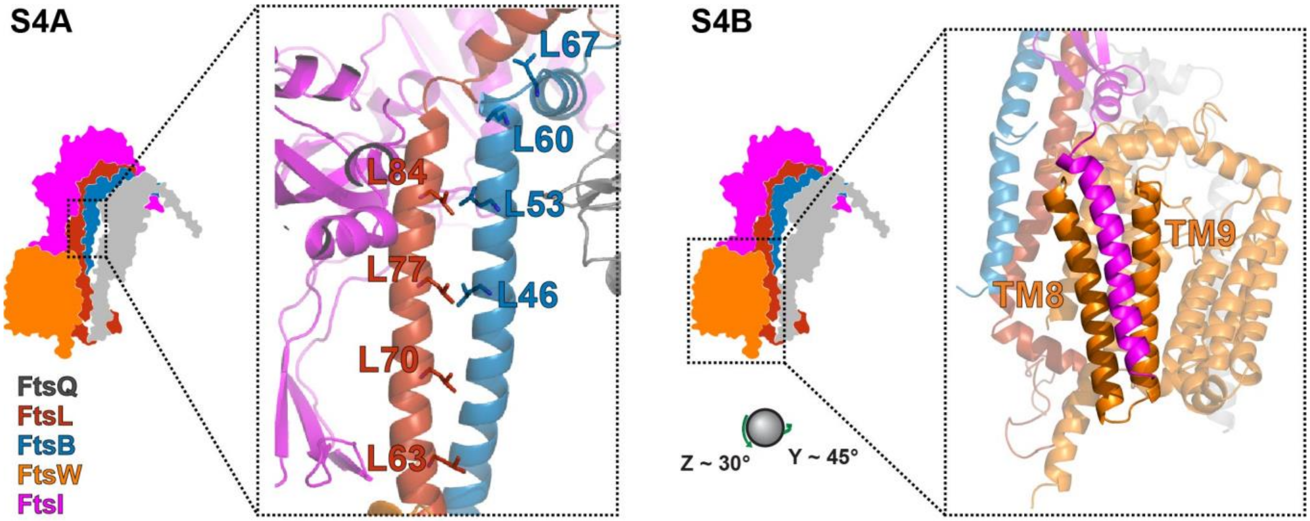


**Supplementary Figure S4.** Helical packing is important in the FtsQLBWI complex. **(A)** A previously predicted leucine-zipper-like motif ($FtsL^{L63, L70, L77, L84}$ and $FtsB^{L46, L53, L60, L67}$) between FtsL and FtsB was predicted by AF2 and persisted during MD [10]. **(B)** Packing between FtsW transmembrane (TM) helices H8 and H9 and the FtsI TM helix is similar to that observed in RodA-PBP2 [11].

### Supplementary Figure S5

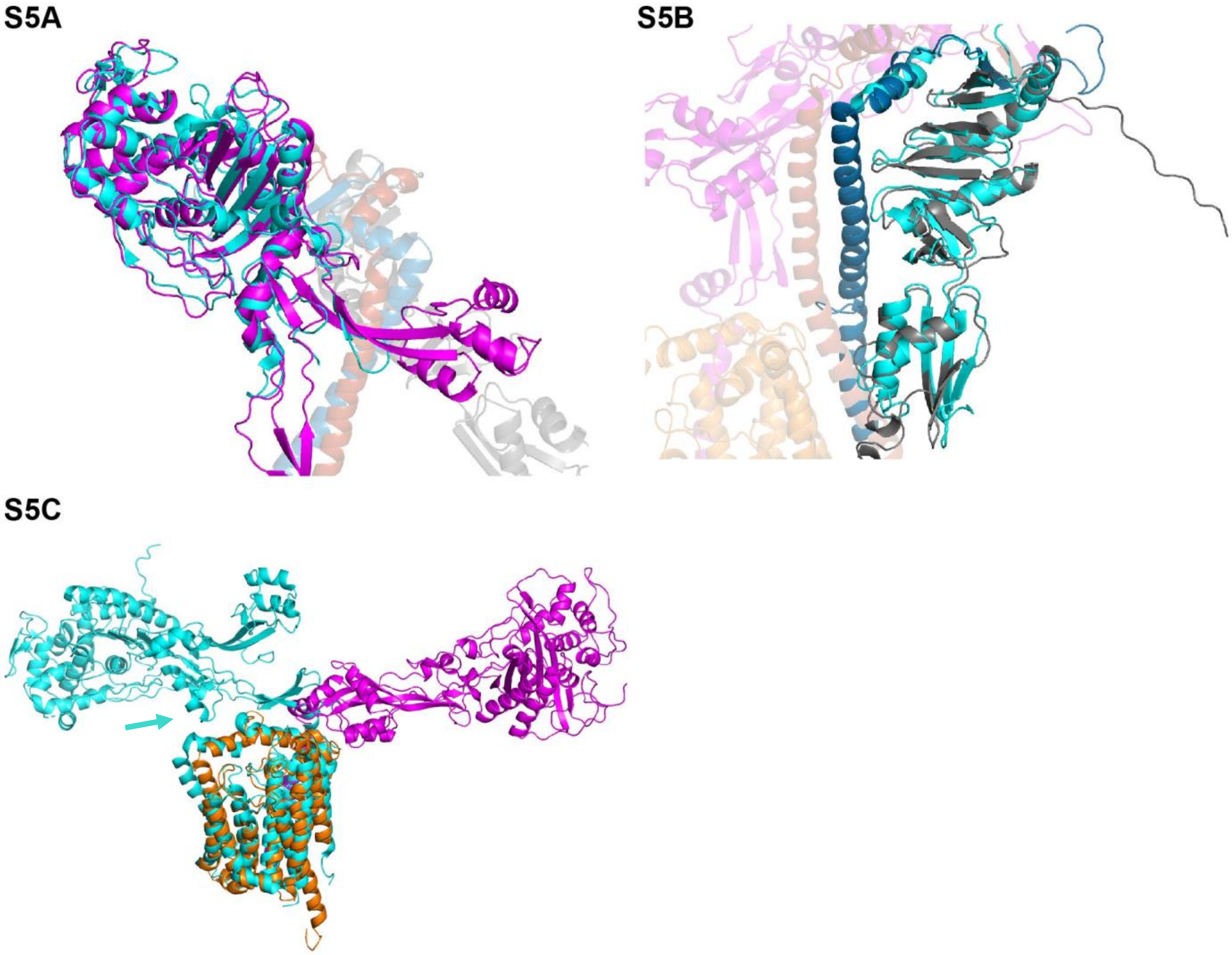


**Supplementary Figure S5.** Comparison between conformers extracted from simulation and existing atomic structures. **(A)** The FtsI TPase domain (magenta) after 1-µs MD is overlaid with crystal structure 7ONO (cyan; residues 89-163 and 204-227 replaced with tri-glycine linkers) of *E. coli* FtsI [12]. **(B)** FtsQ (gray) and FtsB (blue) after 1-µs MD overlaid with the crystal structure (cyan) of FtsQ and a portion of FtsB (5Z2W) from *E. coli* [13]. **(C)** Overlay of FtsWI (orange and magenta) after 1-µs MD with the RodA crystal structure (cyan) 6PL5 from *T. thermophilus* [11] . Note that the periplasmic domains of FtsI in the model (magenta) rotates to the opposite direction of PBP2 in crystal structure (cyan). We note that FtsI lacks nine residues in a loop of PBP2 that mediates PBP2 interaction with RodA ECL4 (cyan arrow).

### Supplementary Figure S6

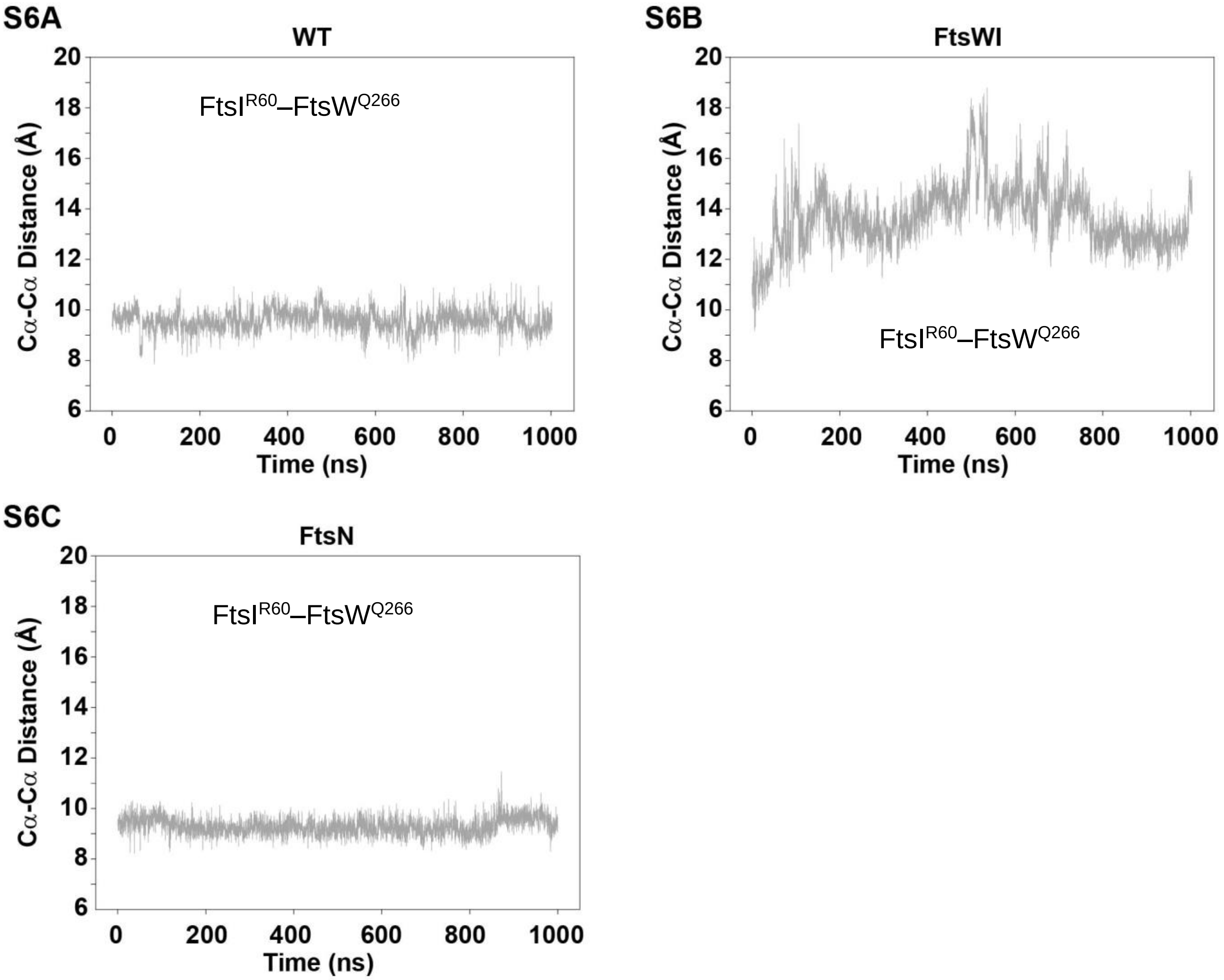


**Supplementary Figure S6.** Dynamics of Cα-Cα distances of FtsI$^{R60}$–FtsW$^{Q266}$ measured from MD trajectories of different complexes show that the predicted hydrogen-bonding interaction between the residues that is frequent in FtsQLBWI (**A**, **Fig. 1E**, **Supplementary Data 1**) is quickly broken by disruption of the Pivot region in FtsWI (**B**), and is maintained in the presence of FtsN (**C**).

### Supplementary Figure S7

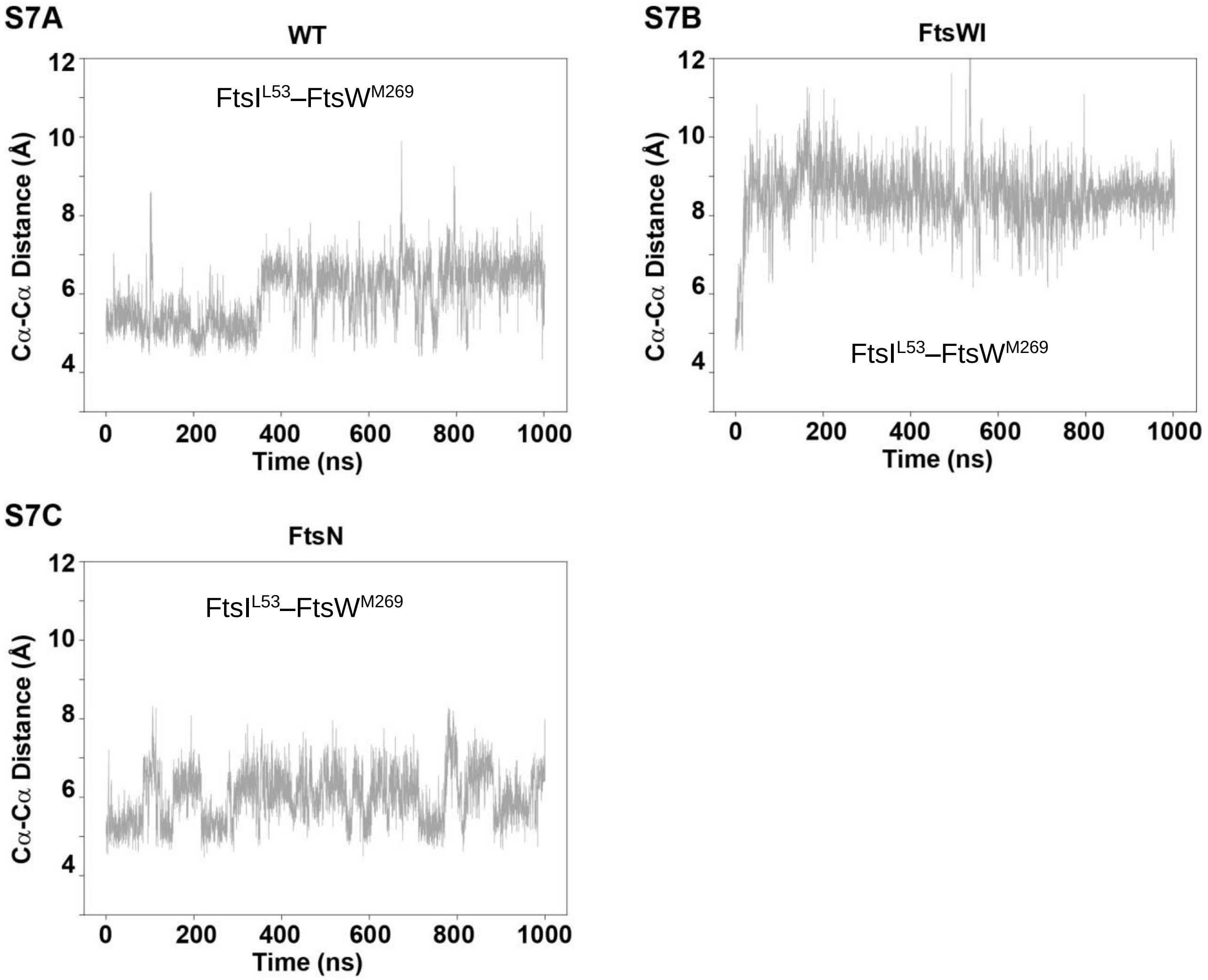


**Supplementary Figure S7.** Dynamics of Cα-Cα distances of $FtsI^{L53}$–$FtsW^{M269}$ measured from MD trajectories of different complexes show that hydrophobic contact between this pair of residues (**Fig. 1E**) is maintained in WT (FtsQLBWI, **A**), rapidly broken by disruption of the Pivot region in FtsWI (**B**), and maintained in the presence of FtsN (**C**).

### Supplementary Figure S8

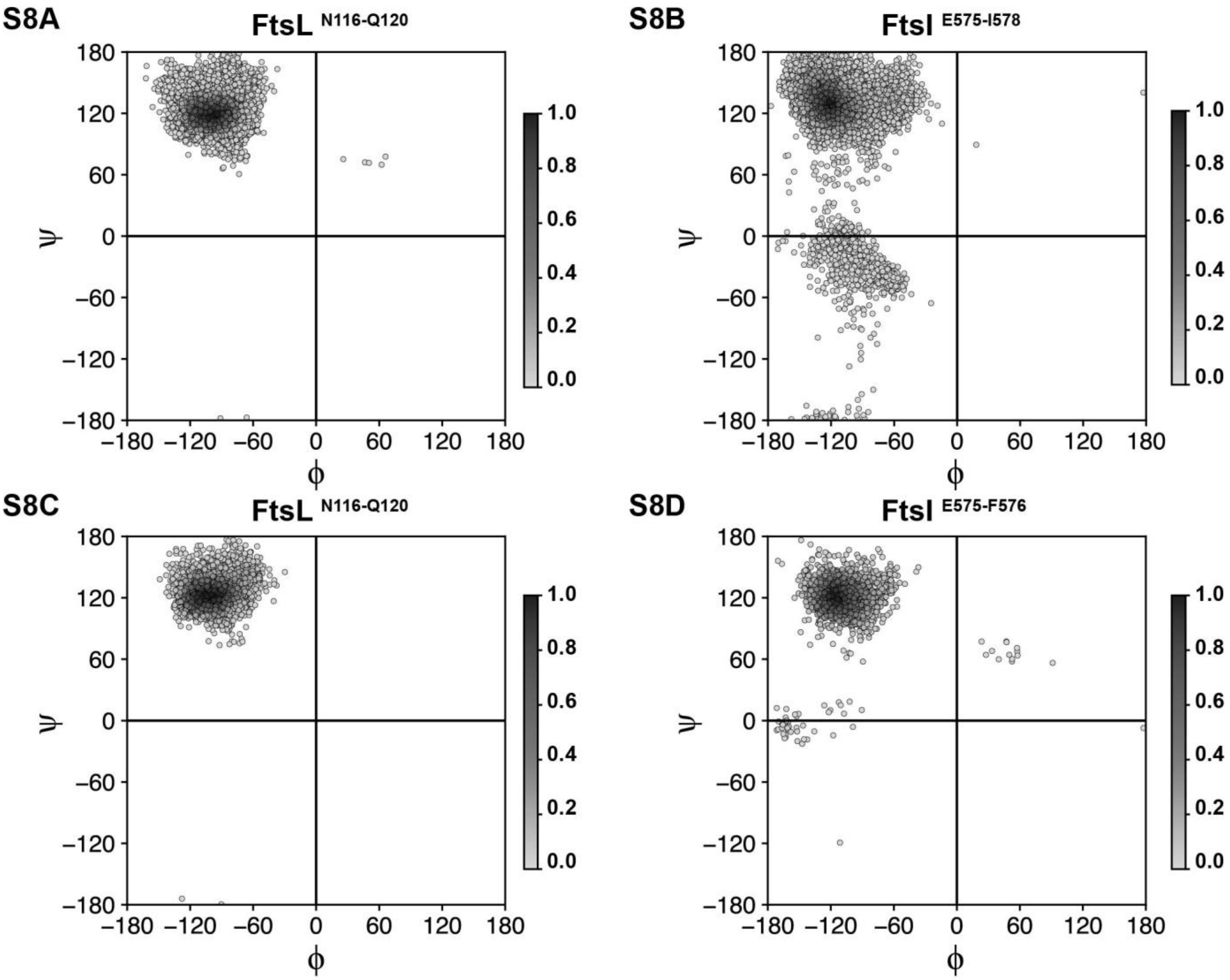


**Supplementary Figure S8.** Dihedral angle plots for residues of FtsL and FtsI C-termini indicate a stable β-sheet. Ramachandran plots colored by density for FtsL residues N116–Q120 **(A)** and FtsI residues E575–I578 **(B)** from the 1-μs FtsQLBWI WT trajectory, and for FtsL residues N116–Q120 **(C)** and FtsI residues E575–I576 **(D)** from the 200-ns FtsI C-terminal truncated trajectory, show stable β-strand conformation at the top left quadrant.

### Supplementary Figure S9

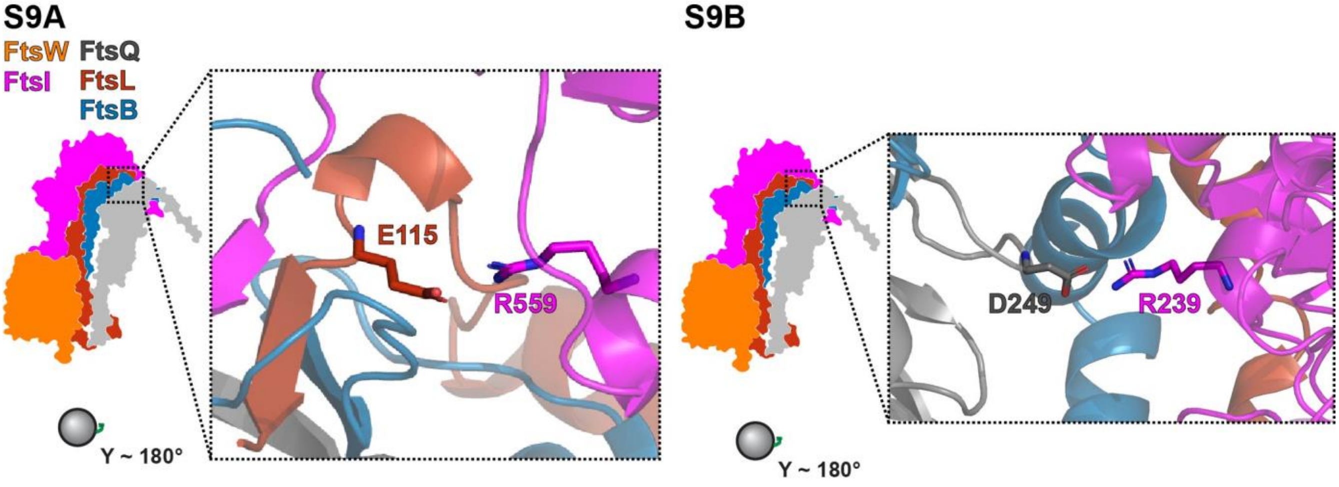


**Supplementary Figure S9.** Hydrogen bonding between $FtsI^{R559}$ and $FtsL^{E115}$ (**A**) and between $FtsI^{R239}$ and $FtsQ^{D249}$ **(B)** is present at the Truss region to tether FtsI to FtsL and FtsQ respectively.

### Supplementary Figure S10

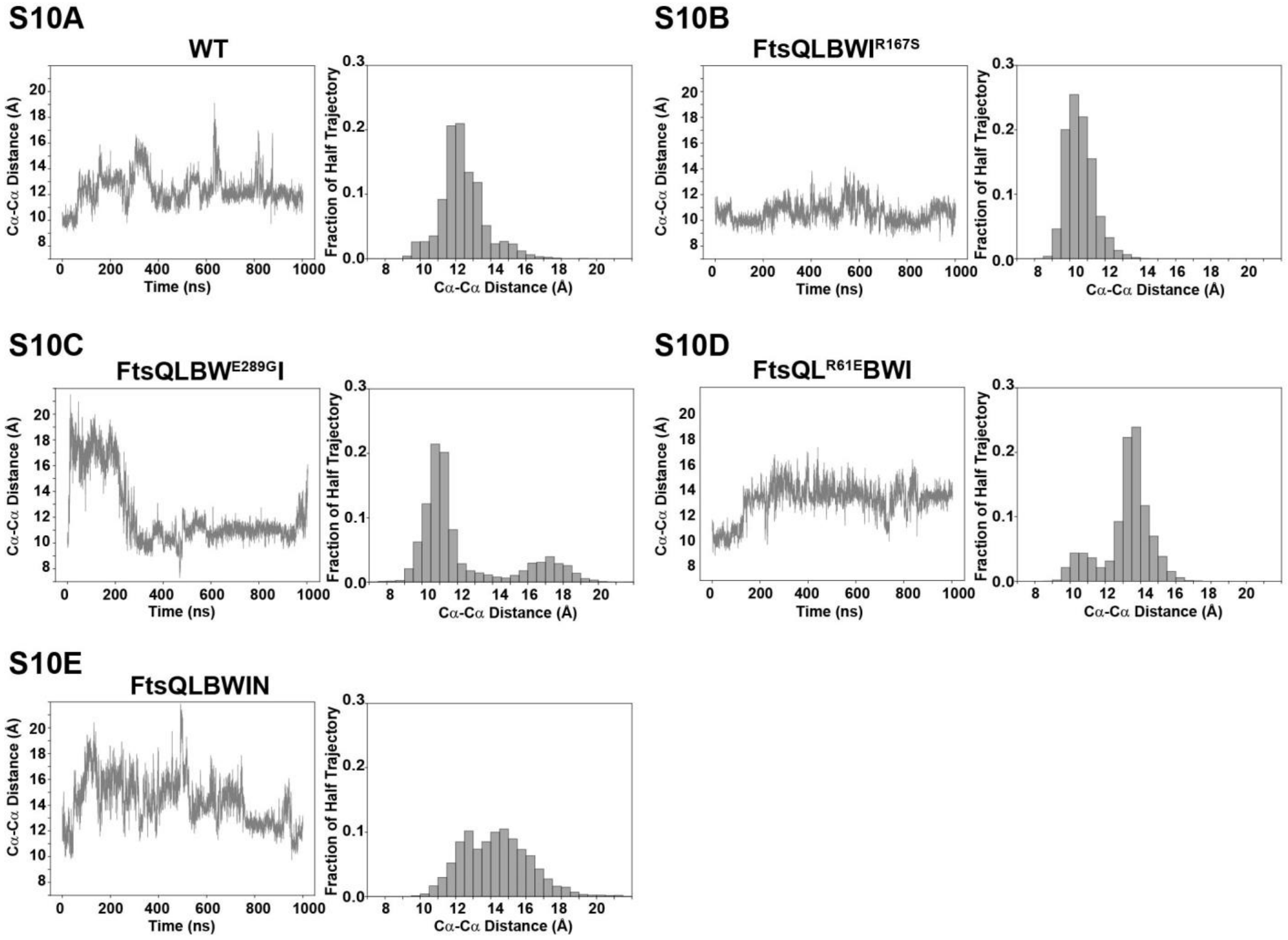


**Supplementary Figure S10.** C$\alpha$-C$\alpha$ distance time traces and corresponding histograms for contact pair FtsI$^{R559}$ and FtsL$^{E115}$ for each 1-μs simulation system show that the contact between the pair fluctuates in different complexes: **(A)** FtsQLBWI (WT), **(B)** FtsQLBWI$^{R167S}$ (R167S), **(C)** FtsQLBW$^{E289G}$I (E289G), (**D**) FtsQL$^{R61E}$BWI (R61E), and **(E)** FtsQLBWIN (FtsN).

### Supplementary Figure S11

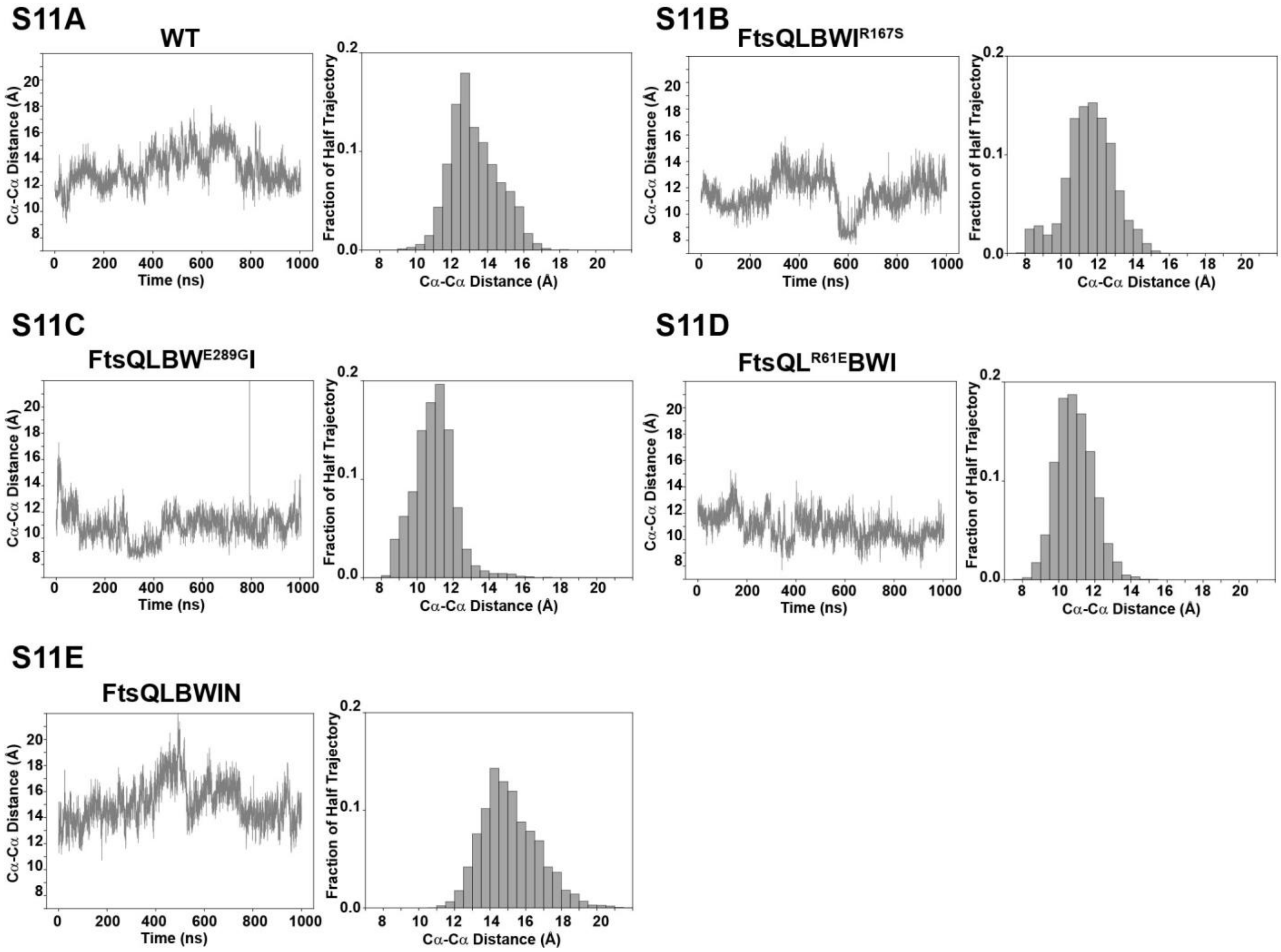


**Supplementary Figure S11.** Cα-Cα distance time traces and corresponding histograms for contact pair FtsI$^{R239}$ and FtsQ$^{D249}$ for each 1-μs simulation system show that the contact between the pair fluctuates in different complexes: **(A)** FtsQLBWI (WT), **(B)** FtsQLBWI$^{R167S}$ (R167S), **(C)** FtsQLBW$^{E289G}$I (E289G), (D) FtsQL$^{R61E}$BWI and **(E)** FtsQLBWIN (FtsN).

### Supplementary Figure S12

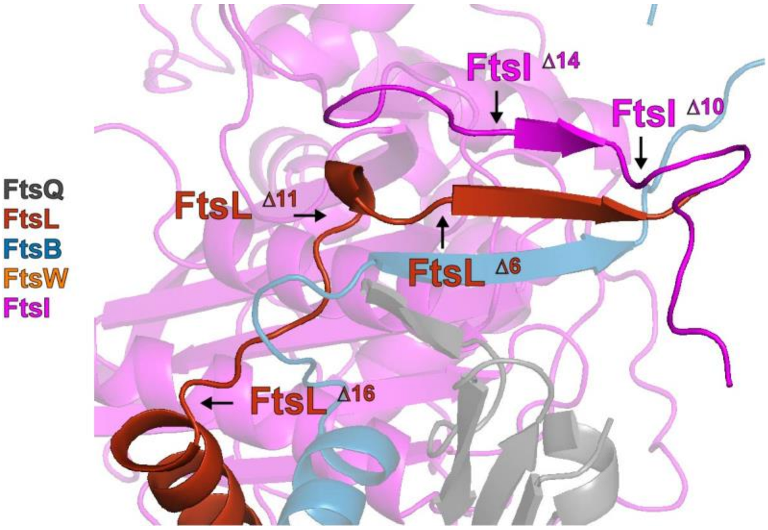


**Supplementary Figure S12.** Schematic of FtsL and FtsI C-terminal truncations. Each arrow points to the position of a truncation mutant. All three FtsL C-terminal truncations (FtsL$^{\Delta6}$, FtsL$^{\Delta11}$, and FtsL$^{\Delta16}$) lack the C-terminal β-strand, and FtsL$^{\Delta11}$ and FtsL$^{\Delta16}$ additionally lack residues such as FtsL$^{E115}$ that interacts with FtsI. FtsI$^{\Delta10}$ only has the disordered C-terminus after the predicted β-strand deleted, and FtsI$^{\Delta14}$ has both the β-strand and the C-terminus deleted. Reported post-translational cleavage of FtsI between FtsI$^{I578\text{-}S588}$ would be FtsI$^{\Delta11}$ following this nomenclature, removing one residue with a hydrophobic sidechain from the predicted β-strand that interacts with FtsL.

### Supplementary Figure S13

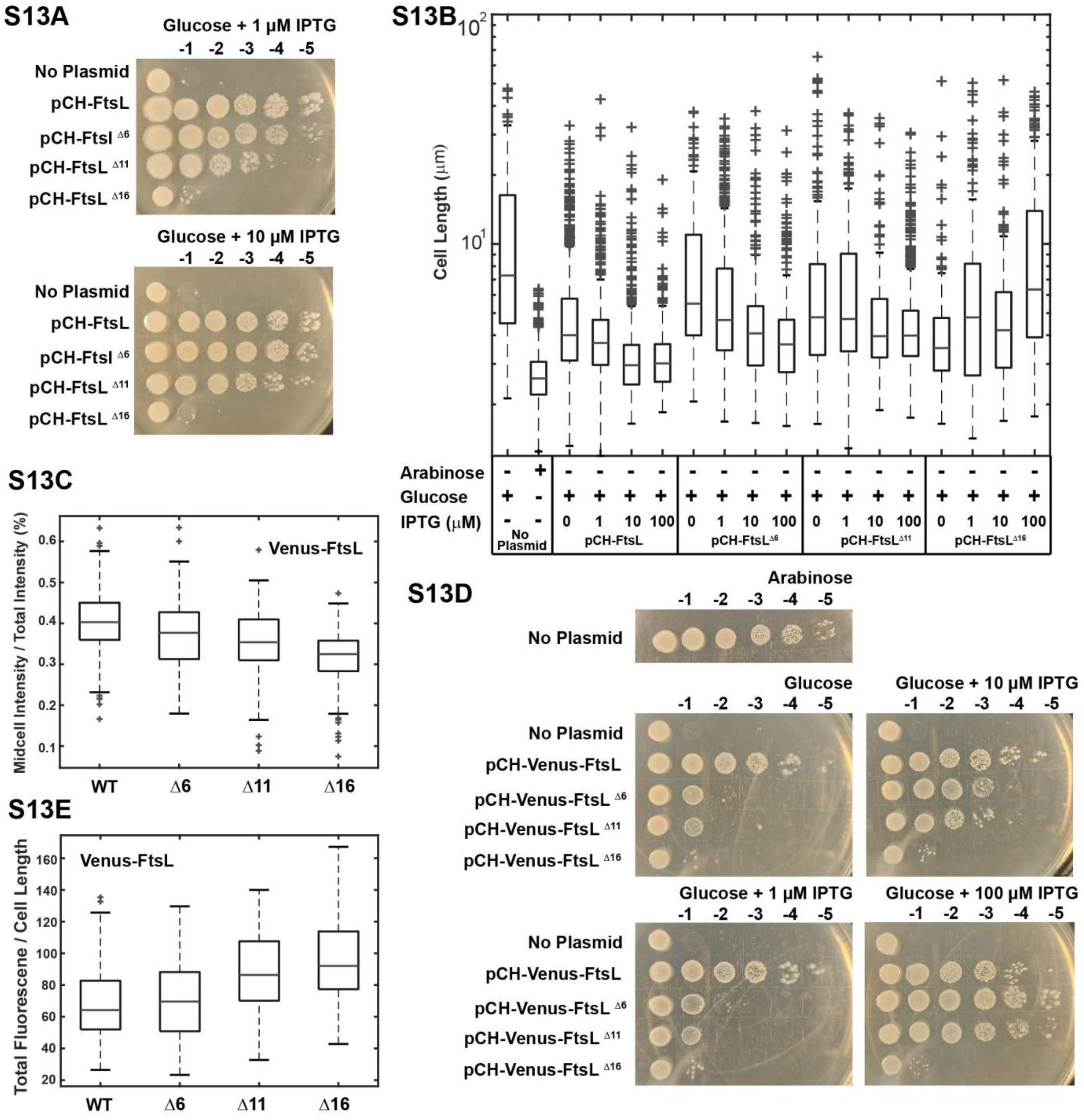


**Supplementary Figure S13.** Functional characterization of FtsL C-terminal truncation mutants. **(A)** Spot dilution complementation test of FtsL truncation mutants. *E. coli* cells depleted of chromosomal wild-type FtsL in which FtsL is expressed from the $P_{BAD}$ promoter (strain MDG279, **Table S2**) failed to complement in the presence of glucose (No Plasmid, first row of both panels). The same depletion strain (MDG279) expressing wild-type FtsL (pCH-FtsL, or pBMB064), $FtsL^{\Delta 6}$ (pCH-$FtsL^{\Delta 6}$, or pBMB065) and $FtsL^{\Delta 11}$ (pCH-$FtsL^{\Delta 11}$, or pBMB066, **Table S1**)

from a *lac* promoter on plasmids complemented the depletion with induction by glucose at both 1 μM and 10 μM IPTG (middle rows of both panels). Expression of FtsL$^{\Delta16}$ was unable to complement at both induction levels (bottom rows of the bottom panel). **(B)** Cell length quantification of *E. coli* cells depleted of wild-type FtsL (No Plasmid) and expressing FtsL WT, FtsL$^{\Delta6}$, FtsL$^{\Delta11}$ and FtsL$^{\Delta16}$ showed that none of the FtsL truncation mutants had WT-like cell lengths even at the highest IPTG induction level (100 μM). Boxplot of cell lengths. Boxes represent interquartile range (IQR) with central mark indicating the median cell length. Boundaries of whiskers based on 1.5 IQR. Outliers shown as +. **(C)** Fluorescence images were quantified to measure the fraction of fluorescence intensity at midcell. The same depletion strain (MDG279) expressing wild-type Venus-FtsL (pCH-Venus-FtsL, or pBMB069), Venus-FtsL$^{\Delta6}$ (pCH-Venus-FtsL$^{\Delta6}$, or pBMB070), Venus-FtsL$^{\Delta11}$ (pCH-Venus-FtsL$^{\Delta11}$, or pBMB071), Venus-FtsL$^{\Delta16}$ (pCH-Venus-FtsL$^{\Delta11}$, or pBMB072, **Table S1**) from a *lac* promoter on plasmids in the presence of glucose and 100 μM IPTG show progressively reduced midcell localization percentage of FtsL truncation mutants. Boxplots as described in (**S13B**) showing midcell fluorescence ratio. **(D)** Complementation test of mVenus-FtsL truncation mutants at different induction levels showed similar result as that in (**A)**: Venus fusions of FtsL$^{\Delta6}$, and FtsL$^{\Delta11}$ complemented the depletion strain at the highest induction level while FtsL$^{\Delta16}$ did not. (**E**) Total cellular fluorescence level normalized against cell length for mVenus-FtsL expressing cells at an induction level of 100 μM IPTG showed that there was not a significant difference in mVenus-FtsL expression levels across the mutants. Boxplots as described in (**S13B**) showing total fluorescence vs cell length.

### Supplementary Figure S14

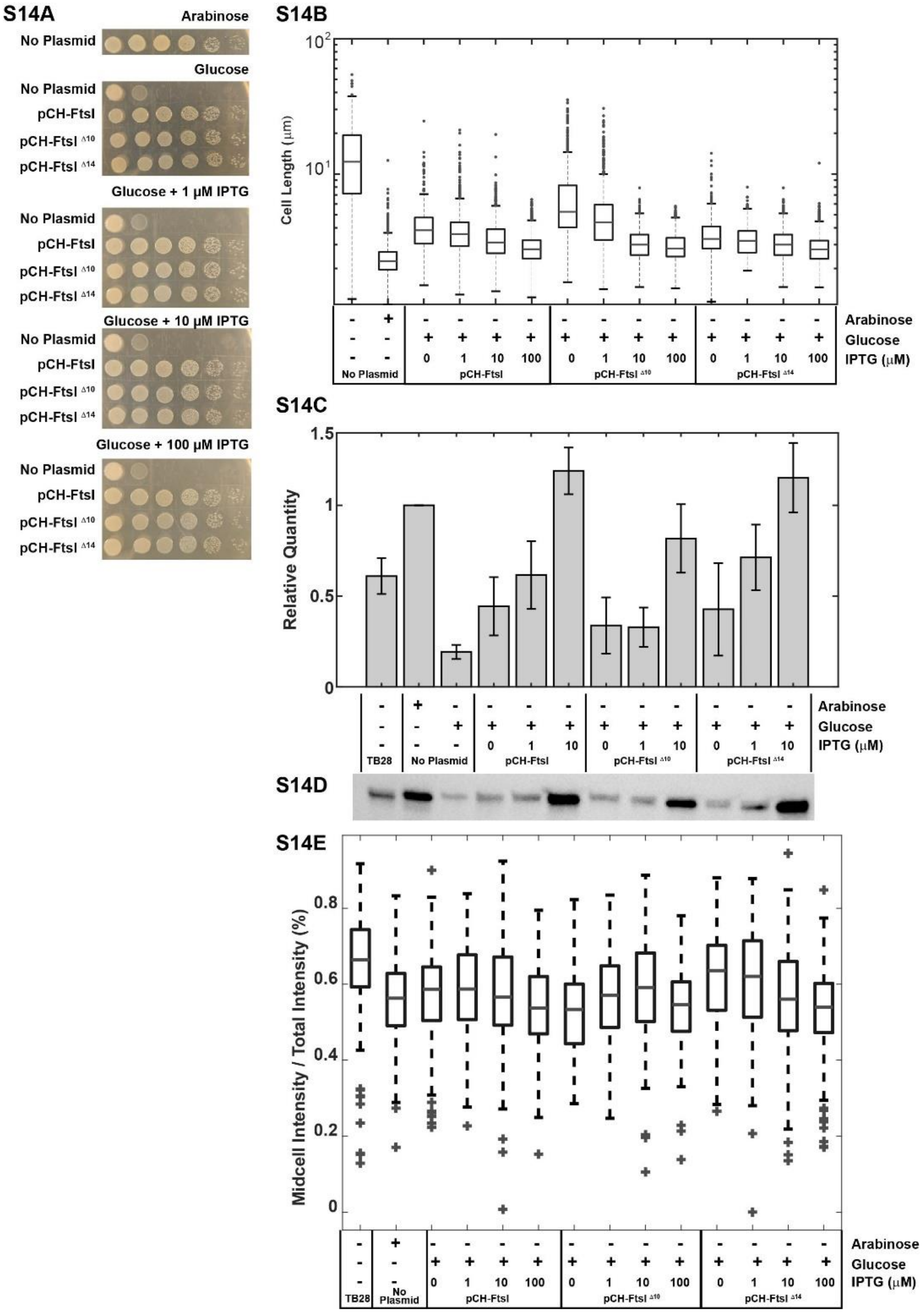


**Supplementary Figure S14. (A)** Spot dilution complementation test of FtsI truncation mutants. *E. coli* cells depleted of chromosomal wild-type FtsI but contain FtsI expressed from the $P_{BAD}$ promoter (strain EC812, **Table S2**) complemented in the presence of arabinose (top panel), but failed in the presence of glucose (No Plasmid, first row of the second panel). The same depletion strain (EC812) expressing wild-type FtsI (pCH-FtsI, or pBMB090), $FtsI^{\Delta 10}$ (pCH-$FtsI^{\Delta 10}$, or pBMB092), and $FtsI^{\Delta 14}$ (pCH-$FtsI^{\Delta 14}$, or pBMB093, **Table S1**) from a *lac* promoter on plasmids complemented the depletion in the presence of glucose at both no induction and 1 μM, 10 μM, and 100 μM IPTG conditions (middle rows of lower panels). **(B)** Cell length quantification for *E. coli* cells depleted of wild-type FtsI and expressing wild-type FtsI, FtsI$^{\Delta10}$, or FtsI$^{\Delta14}$ show that, at low induction levels (0 and 1 μM IPTG), FtsI$^{\Delta10}$ cells are longer than WT and FtsI$^{\Delta14}$ cells. Boxplot of cell lengths. Boxes represent interquartile range (IQR) with central mark indicating the median cell length. Boundaries of whiskers based on 1.5 IQR. Outliers shown as +. **(C)** Quantification of protein expression levels normalized to the parent strain, EC812, expressing FtsI from the $P_{BAD}$ promoter with induction by arabinose. TB28 represents a wildtype strain with endogenous expression levels of FtsI. Truncation to FtsI$^{\Delta10}$ results in lower FtsI expression levels than expression of wild-type FtsI or FtsI$^{\Delta14}$. **(D)** A representative anti-FtsI western blot used for quantifying FtsI expression levels. **(E)** Anti-FtsI immunofluorescence images were quantified to measure the fraction of fluorescence intensity at midcell. Boxplots as described in (**S14B**) showing midcell fluorescence ratio.

### Supplementary Figure S15

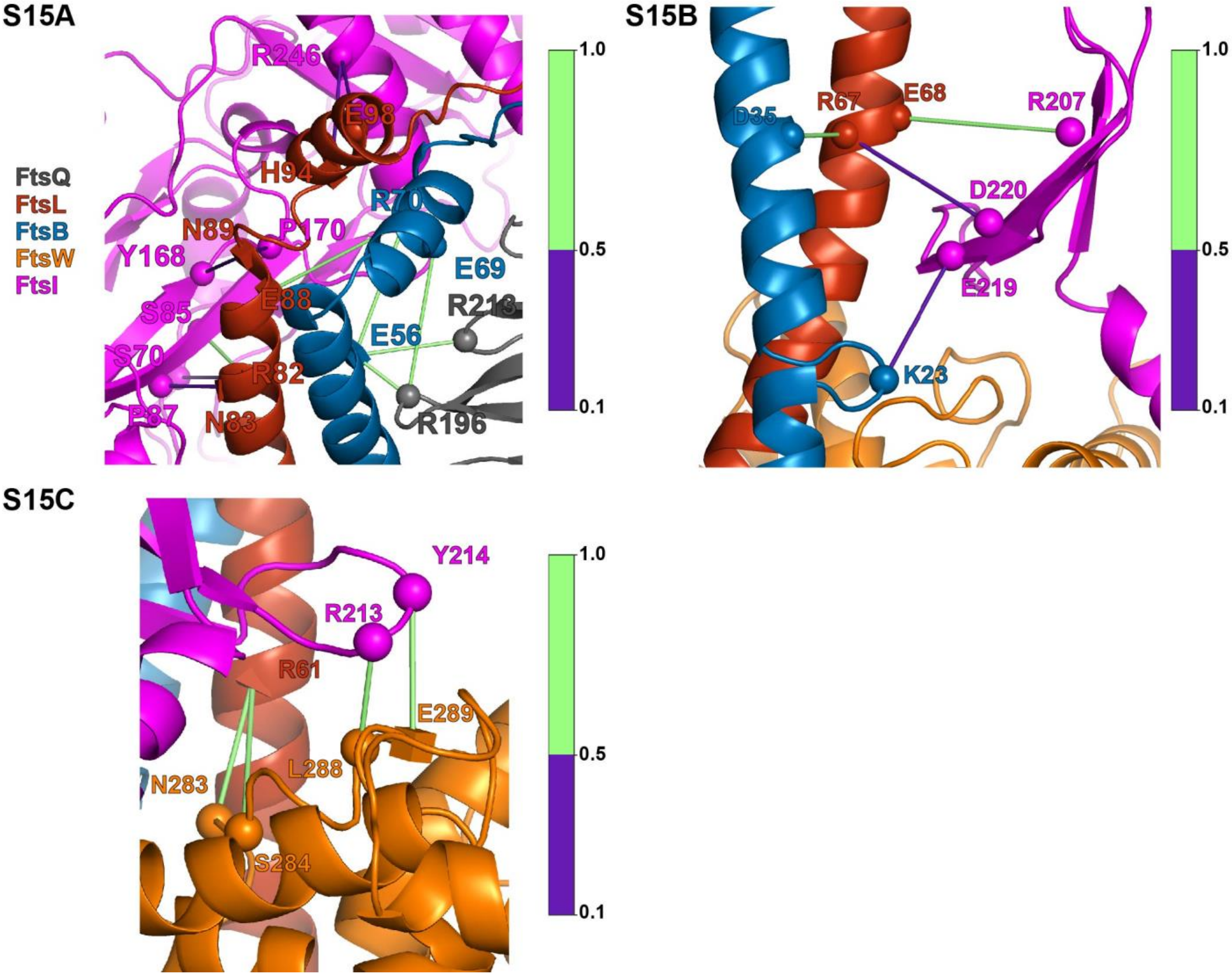


**Supplementary Figure S15.** Depth cued versions of main text figures showing hydrogen-bonding network. Spheres indicate residues involved in hydrogen bonding. Superfission mutant residues are depicted as squares. Dominant negative mutant residues depicted as tetrahedrons. Hydrogen-bond frequencies during the last 500 ns of FtsQLBWI simulation are indicated by the colorbar. **(A)** Corresponds to **Figure 3A**. A hydrogen bonding network extends from FtsQLB to FtsI in the CCD interface of the Hub region. **(B)** Corresponds to **Figure 4A**. Hydrogen bonding between the anchor domain of FtsI and the helices of FtsLB fix the position of FtsI anchor loop over FtsW ECL4. **(C)** Corresponds to **Figure 4B**. Hydrogen bonding between $\text{FtsL}^{R61}$ positions the ECL4 of FtsW under the anchor domain of FtsI. $\text{FtsI}^{Y214}$ hydrogen bonds with $\text{FtsW}^{E289}$ positioning FtsW ECL4 under the FtsI anchor.

### Supplementary Figure S16

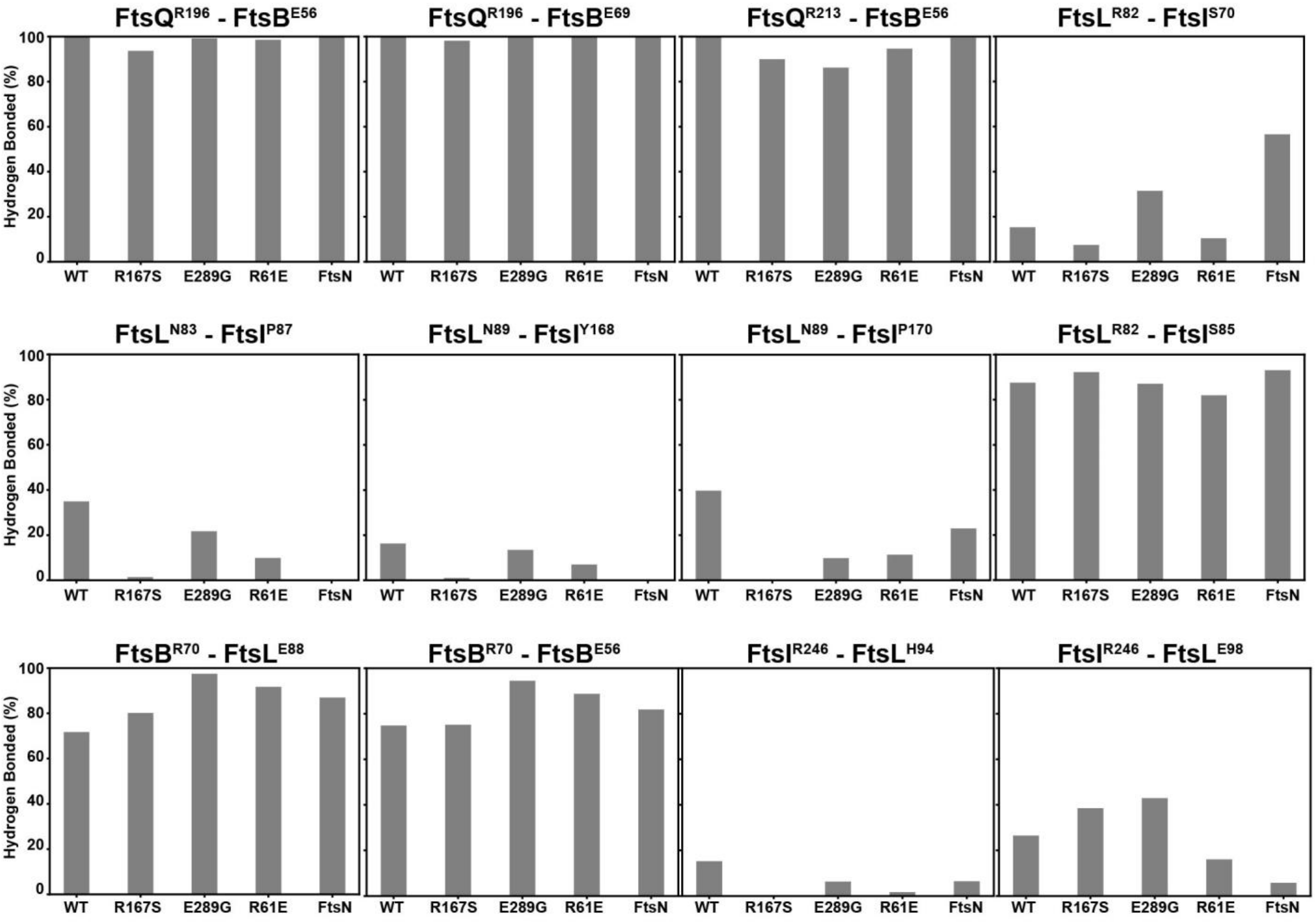


**Supplementary Figure S16.** Hydrogen bonding frequency is plotted for the network of interactions at the Truss region, related to **Figure 3A**. On each bar graph the hydrogen bond frequency throughout the last 500 ns of each simulation systems is plotted: FtsQLBWI (WT), FtsQLBWI$^{R167S}$ (R167S), FtsQLBW$^{E289G}$I, FtsQL$^{R61E}$ (R61E), and FtsQLBWIN (FtsN).

### Supplementary Figure S17

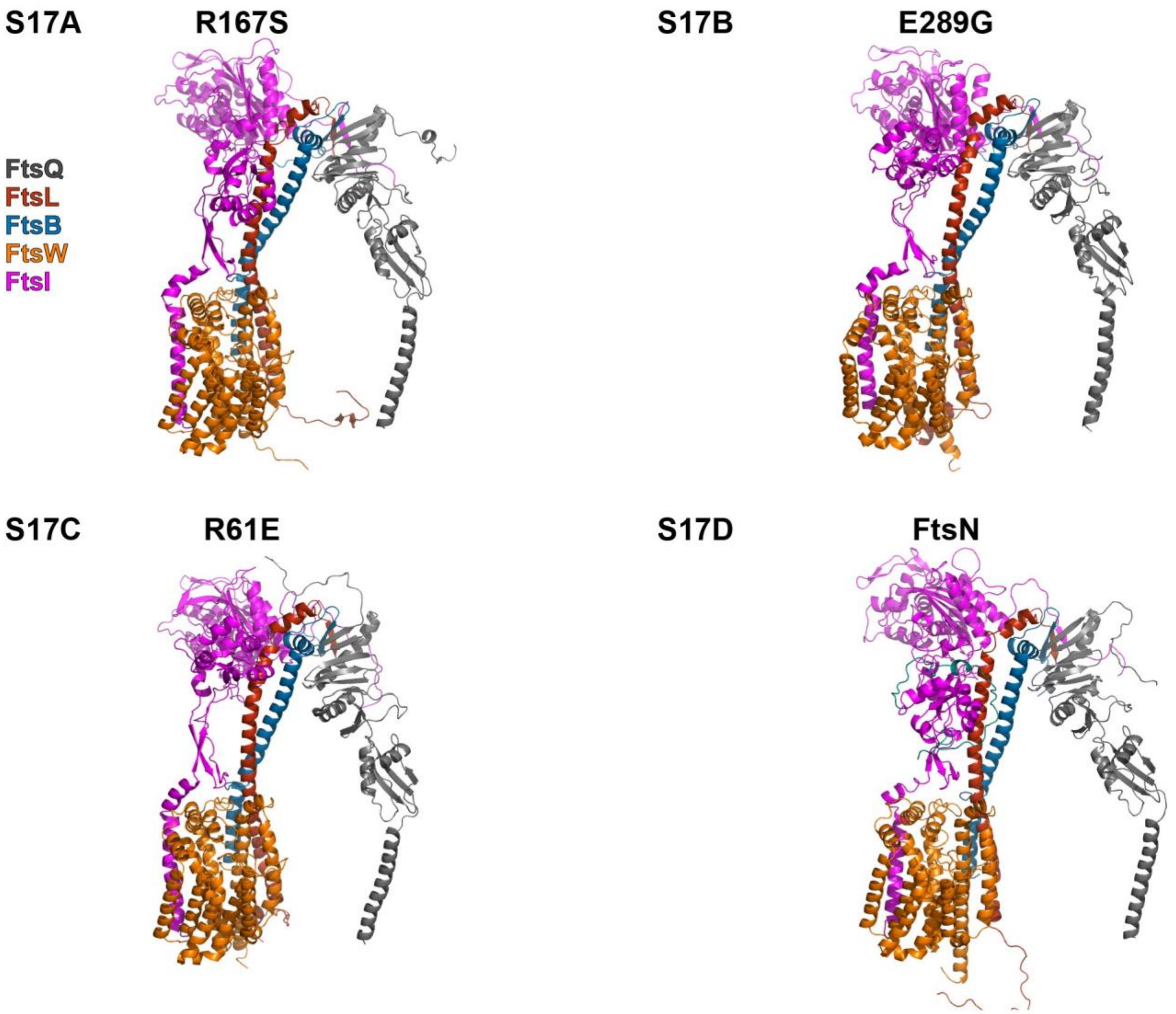


**Supplementary Figure S17.** Conformers after 1-µs MD for each of the following simulation systems: **(A**) FtsQLBWI$^{R167S}$ (R167S), **(B)** FtsQLBW$^{E289G}$I (E289G) (**C**) FtsQL$^{R61E}$BWI (R61E), and **(D)** FtsQLBWIN (FtsN). Here, all conformers are aligned to the orientation of FtsW and PDB files are available for download (**Supplementary Data 1**).

### Supplementary Figure S18

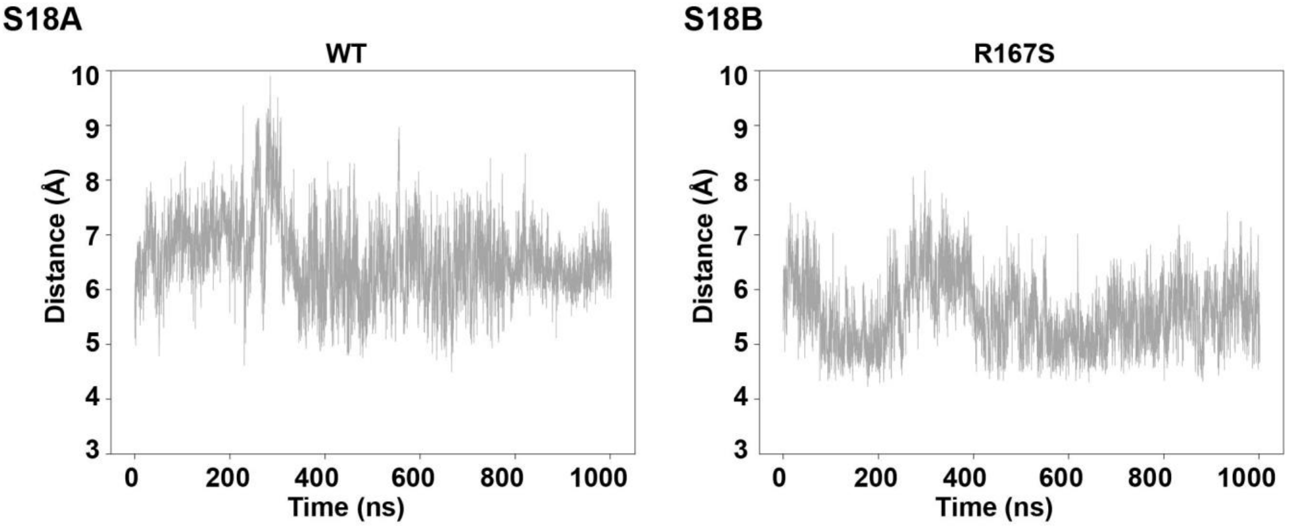


**Supplementary Figure S18.** $FtsI^{V84}$–$FtsL^{I85}$ distance time traces for interactions highlighted in main-text histograms. Distances between sidechain centers of geometry were plotted over time for the entire 1-µs MD trajectory for **(A)** FtsQLBWI (WT) and **(B)** $FtsQLBWI^{R167S}$ (R167S).

### Supplementary Figure S19

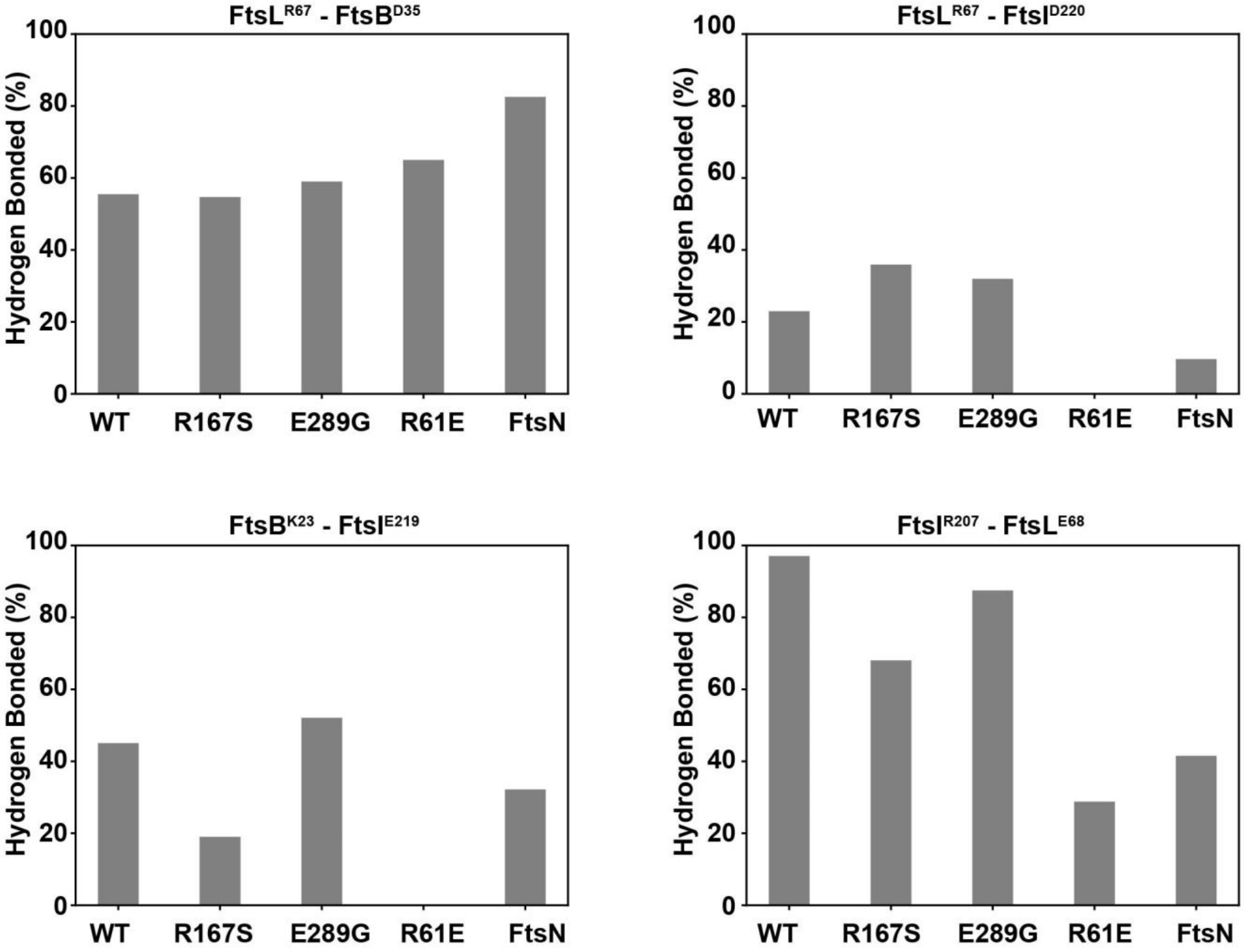


**Supplementary Figure S19.** Hydrogen bonding frequencies in the network of interactions at the Lid region between $\text{FtsL}^{R67}$-$\text{FtsB}^{D35}$, $\text{FtsL}^{R67}$-$\text{FtsI}^{D220}$, $\text{FtsB}^{K23}$-$\text{FtsI}^{E219}$, and $\text{FtsI}^{R207}$- $\text{FtsL}^{E68}$ during the final 500 ns of WT (FtsQLBWI), $\text{FtsQLBWI}^{R167S}$ (R167S), $\text{FtsQLBW}^{E289G}\text{I}$, $\text{FtsQL}^{R61E}$ (R61E), and FtsQLBWIN (FtsN) simulations. The frequencies of all hydrogen bonding were higher in WT and active complexes (R167S, E289G and +FtsN) than that in the dominant negative mutant R61E complex, except for $\text{FtsL}^{R67}$-$\text{FtsB}^{D35}$.

### Supplementary Figure S20

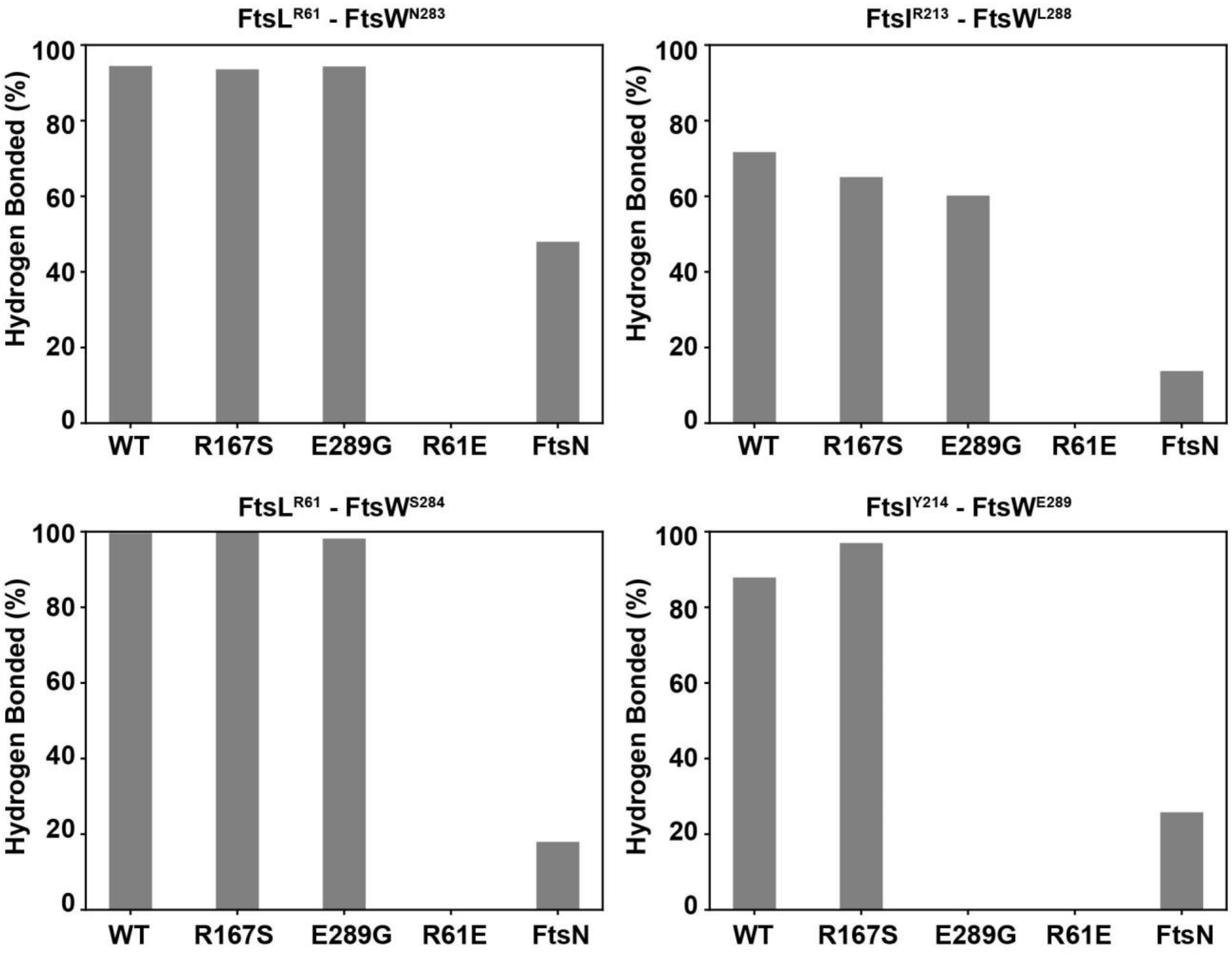


**Supplementary Figure S20.** Hydrogen bonding frequency is plotted for the network of interactions at the Lid region, related to **Fig. 4B.** The hydrogen bond frequency throughout the last 500 ns of each simulation is plotted for each complex: FtsQLBWI (WT), $FtsQLBWI^{R167S}$ (R167S), $FtsQLBW^{E289G}I$, $FtsQL^{R61E}$ (R61E), and FtsQLBWIN (FtsN).

### Supplementary Figure S21

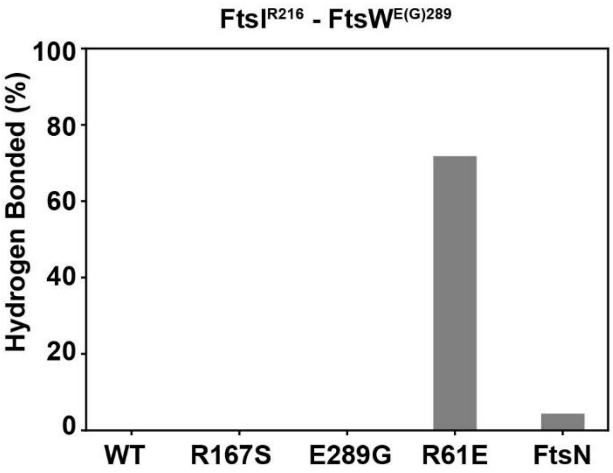


**Supplementary Figure S21.** Hydrogen bonding frequency of $FtsI^{R216}$ and $FtsW^{E(G)289}$ during the last 500 ns of simulations for FtsQLBWI (WT), $FtsQLBWI^{R167S}$ (R167S), $FtsQLBW^{E289G}I$, $FtsQL^{R61E}$ (R61E), and FtsQLBWIN (FtsN) shows that this interaction is only present in the dominate negative R61E complex.

### Supplementary Figure S22

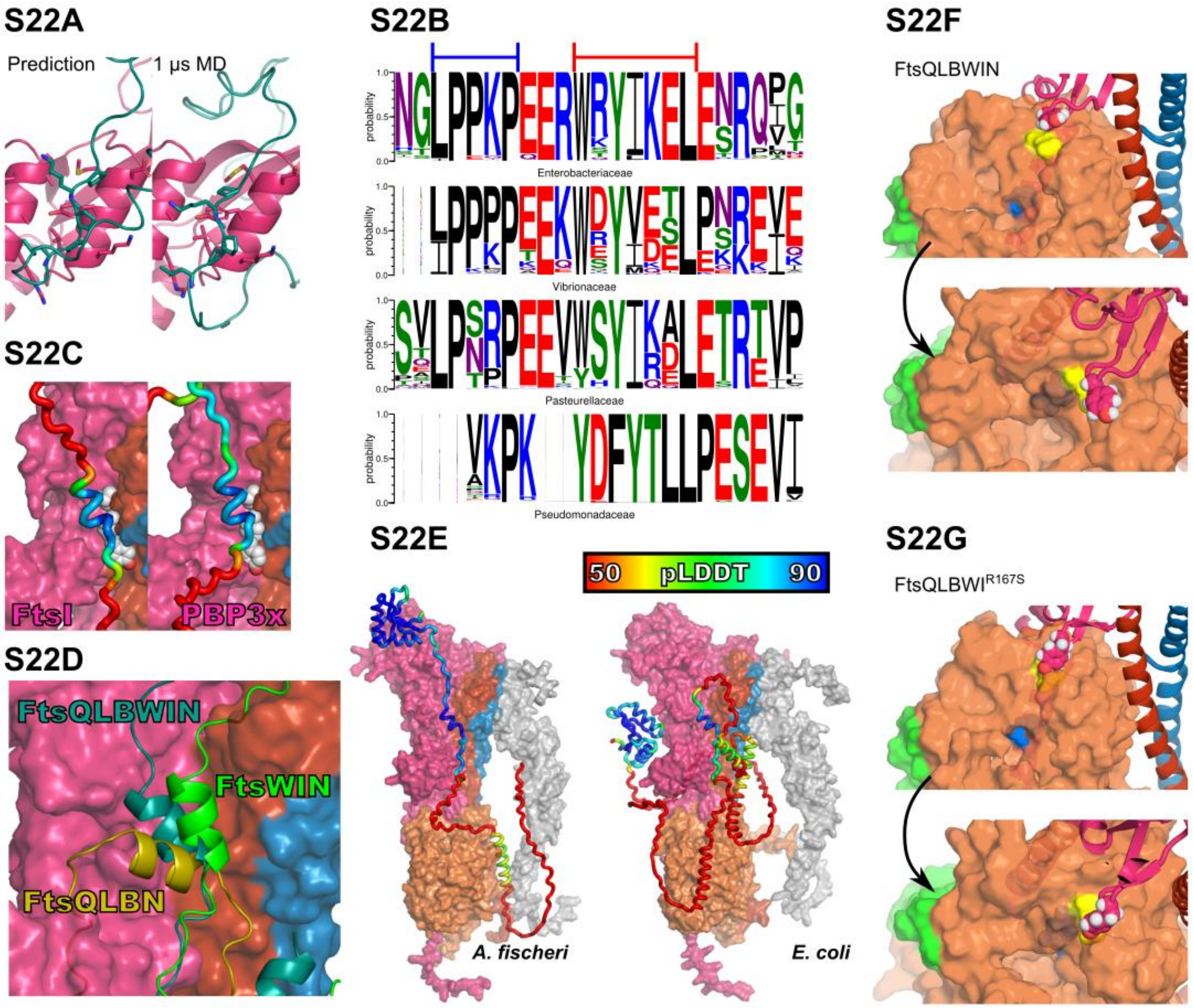


**Supplementary Figure S22.** (**A**) Interactions between residues in the FtsI head domain and FtsN in the structure prediction is compared to interactions after 1-µs MD. Structures are aligned to the FtsI head domain and FtsN$^{75–79}$ is shown together with FtsI residues with atoms within 4 Å of FtsN$^{75–79}$ atoms. (**B**) Sequence logos generated from alignments of regions homologous to *E. coli* FtsN$^{73–95}$ for different γ-proteobacteria families. Column heights indicate probability of aligned residues and column widths indicate likelihood a sequence has no gap at a given position. The WYL motif in *E. coli* FtsN that binds at the AWI interface (red line) is conserved within families. The region in FtsN$^{E}$ with high polyproline-II helix propensity predicted to bind the FtsI head domain (blue line; *e.g.* P, L, R, A, and K [14]) is poorly conserved in *Pseudomonadacea.* (**C**) Complexes for FtsQLBWIN homologues in *Pseudomonas aeruginosa* with both FtsI (left) and PBP3x (right) to observe predicted binding of FtsN$^{68–118}$ (loop, colored by pLDDT). A motif homologous to that in *E. coli* binds the AWI interface between FtsI/PBP3x and FtsL (with the *E. coli* WYL motif substituted by *P. aeruginosa* FtsN$^{Y93}$, FtsN$^{F95}$, and FtsN$^{L99}$ (gray spheres). However, there is no predicted binding to the TPase head domain and very low pLDDT. (**D**) Complexes of *E. coli* FtsN were predicted with FtsQLB (FtsQLBN) and FtsWI (FtsWIN) and compared to the FtsQLBWIN prediction. Predictions were aligned by regions in contact with FtsN in FtsQLBWIN (FtsQLBN: $FtsL^{80–96}$; FtsWIN: $FtsI^{108–285}$) to compare the relative site of predicted FtsN binding. Surface representations show FtsI (magenta), FtsL (red), and FtsB (blue) in FtsQLBWIN and cartoon representation show FtsN from FtsQLBWIN (teal), FtsQLBN (yellow), and FtsWIN (green). (**E**) Predicted structures for FtsQLBWIN with full-length FtsN are shown for *Aliivibrio fischeri* (left) and *E. coli* (right). FtsQLBWI subunits are colored as in other figures and shown in surface representation. FtsN is shown in a tube representation and colored by pLDDT. Although *A. fischeri* has a relatively short linker between $FtsN^{E}$ and the FtsN SPOR domain and appears to have predicted binding to the FtsI TPase domain, few residues have predicted contacts, giving a low pLDDT of 0.14 for this interface. (**F**, **G**) Visualizations from the same points of view shown in **Fig. 4C** are shown for conformers after 1-µs MD for FtsQLBWIN (**F**) and $FtsQLBWI^{R167S}$ (**G**) to illustrate positions of the FtsI anchor-loop containing $FtsI^{Y214}$ (purple) relative to the FtsW catalytic pore (orange), with colors highlighting FtsW TM7 (green), $FtsW^{D297}$ (blue) and $FtsW^{E289}$ (yellow) residues. In both complexes, the anchor-loop is positioned above $FtsW^{E289}$ in ECL4 as observed in FtsQLBWI, but not $FtsQL^{R61E}BWI$ (**Fig. 4C** and **4E**). FtsQLBWIN exhibits a similar change in conformation to $FtsI^{Y214}$ when interaction with $FtsW^{E289}$ is lost to that observed in the $FtsQLBW^{E289G}I$ simulation (**Fig. 4D**)

### Supplementary Figure S23

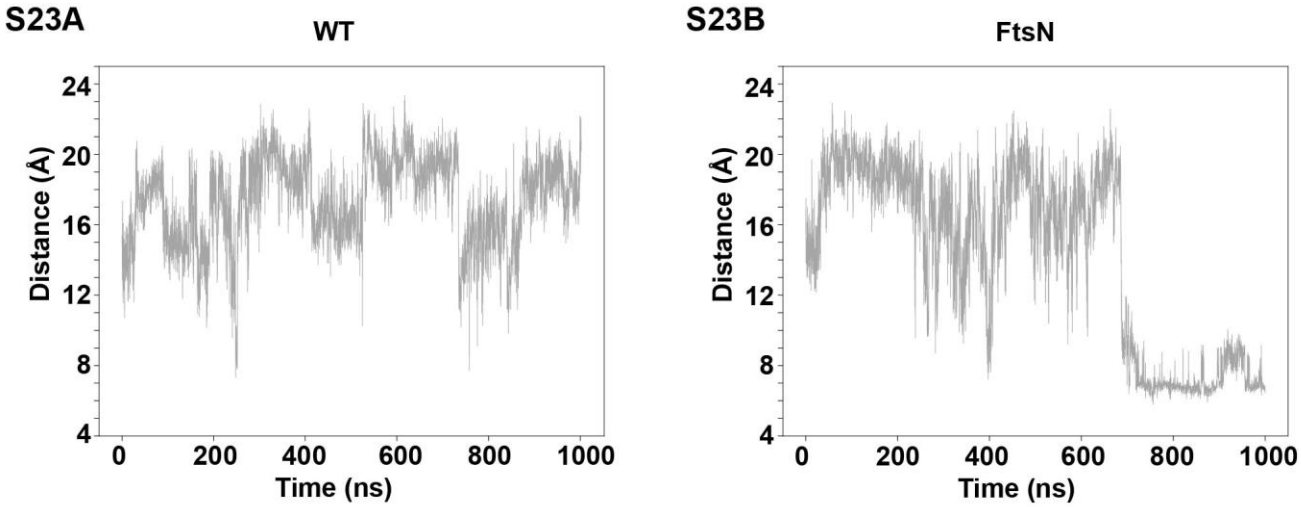


**Supplementary Figure S23.** Time traces of the distances between $FtsL^{R67}$ –$FtsI^{D225}$ sidechain centers of geometry for the entire 1-µs MD trajectory of **(A)** FtsQLBWI (WT) and **(B)** FtsQLBWIN (FtsN) shows that this charged interaction forms in the presence of FtsN in the final third of the 1-µs trajectory.

### Supplementary Figure S24

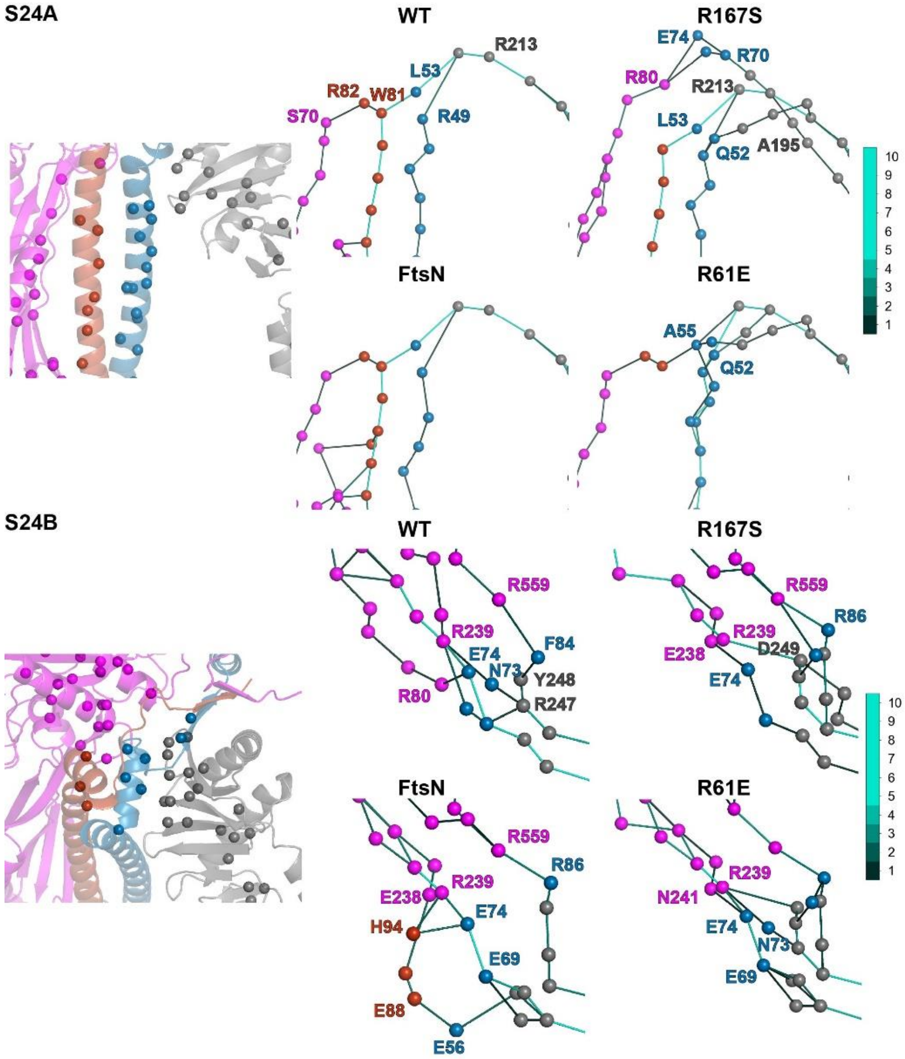


**Supplementary Figure S24.** Long-range interaction pathways showcase allostery in FtsQLBWI complexes. **(A)** Ensembles of optimal paths extending from FtsQ to active-site residue FtsW$^{D297}$ highlight contacts through FtsL, except in the FtsQL$^{R61E}$BWI mutant, that interact through direct FtsB-FtsW contacts. Each sphere represents a residue in the path ensemble. The path ensemble comprises 10 paths, computed for each 50-ns window of the last 500 ns of MD simulation. The color of the line connecting each pair of residues corresponds to the number of paths in the 10-path ensemble in which that pair is present (see colorbar). The paths shown here are the same as in **Fig. 5E** but are from a different perspective to illustrate optimal paths through the Hub region. **(B)** The ensembles of optimal paths from FtsQ to active-site residue FtsI$^{S307}$ is shown to compare paths from FtsQ, through the Hub region, to the active site (not shown in view). We focus here on key residues at the interfaces formed in the hub region. Unique routes of communication were observed across systems, with the path ensemble for FtsN extending through mutation sites FtsB$^{E56}$, FtsL$^{E88}$, and FtsL$^{H94}$.

### Supplementary Figure S25

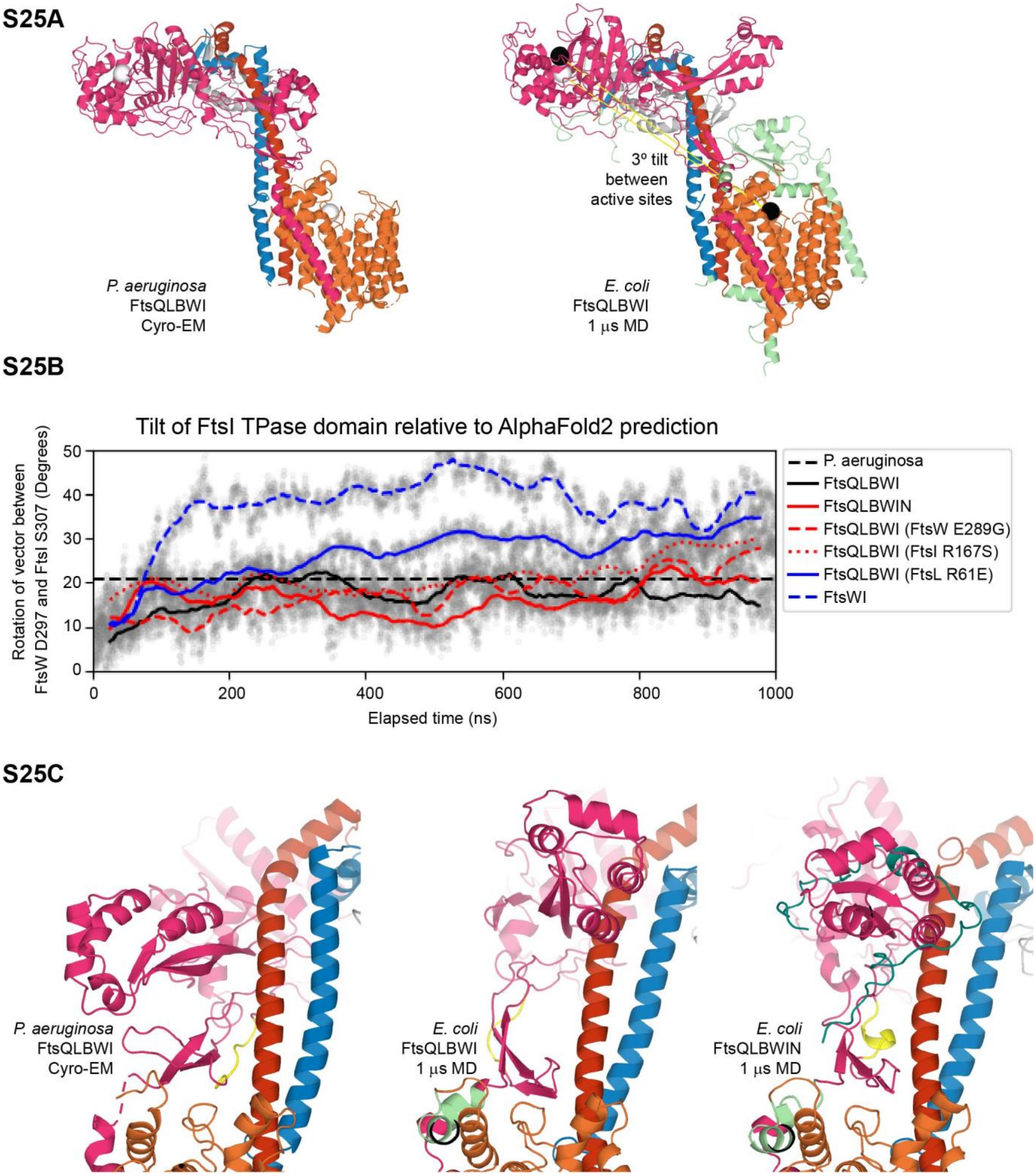


**Supplementary Figure S25.** Comparison of the cryo-EM structure of the *P. aeruginosa* divisome (PDB 8BH1) to the predicted structure and dynamics of the *E. coli* divisome [15] Coloring of divisome subunits in all subfigures: gray, FtsQ; red, FtsL; blue, FtsB; orange, FtsW; magenta, FtsI; cyan, FtsN. (**A**) Left, *P. aeruginosa* cryo-EM structure with $FtsW^{D275}$ and $FtsI^{S294}$ α-carbons shown as white spheres. Right, *E. coli* FtsQLBWI after 1 µs MD after alignment to *P. aeruginosa* FtsW residues. Loops and terminal residues included in MD simulations, but unresolved in the *P. aeruginosa* structure, are shown in pale green. Black spheres show *E. coli* $FtsW^{D297}$ and $FtsI^{S307}$ α-carbons and white spheres indicate positions of equivalent *P. aeruginosa* atoms. The angle formed between the vectors connecting active-site α-carbons is 3 degrees (yellow). (**B**) The tilt of the FtsI TPase domain relative to its initial position in an AF2 prediction of the *E. coli* FtsQLBWI structure is plotted over 1 µs of MD simulation. All trajectories are aligned relative to initial FtsW coordinates. The tilt angle is formed by vectors between $FtsW^{D297}$ and $FtsI^{S307}$ ($FtsW^{D275}$ and $FtsI^{S294}$ for *P. aeruginosa*). The *P. aeruginosa* FtsQLBWI cryo-EM structure exhibits a 21-degree tilt relative to predicted *E. coli* coordinates (black dashed line). Transparent gray circles show angles calculated for individual frames to illustrate dynamics. Other lines show median tilt angles over 50-ns time windows. Refer to the inset legend for trajectory identities. Angles for wild-type FtsQLBWI complexes are colored black, angles with FtsN or SF complexes are colored red, and angles for partial or DN complexes are colored blue. (**C**) Left: the *P. aeruginosa* cryo-EM structure was reported to exhibit direction interaction between the FtsI head and anchor domains and to form an interface between $FtsI^{208–212}$ not observed in an AF2 prediction of the complex; here, residues 209–215 are colored yellow. Middle: neither conformational change is observed for *E. coli* FtsQLBWI after 1 µs MD; here, $FtsI^{223–228}$ (equivalent to *P. aeruginosa* $FtsI^{209–215}$) is colored yellow. Right: following 1 µs MD in an *E. coli* complex including FtsN, $FtsI^{223–228}$ (yellow) shift to form contacts with FtsL and FtsN bridges the FtsI head and anchor domains. Views are rotated 90 degrees relative to those in **A**.

**Movie S1.** Epifluorescence time-lapse imaging of a single Halo-FtsB molecule in M9-glucose, corresponding to Figure 1A (top). Scale bar, 0.5 µm.

**Movie S2.** Epifluorescence time-lapse imaging of a single Halo-FtsB molecule in M9-glucose, corresponding to Figure 1A (bottom). Scale bar, 0.5 µm.

**Movie S3.** One-microsecond simulation of the FtsQLBWI complex.

**Movie S4.** One-microsecond simulation of FtsWI embedded in a lipid bilayer. Phosphorus (dark purple) and oxygen (light purple) atoms of the polar head groups are shown.

**Movie S5.** One microsecond simulation of the FtsQLBWI complex with bound activator FtsN.

### References


1 Buss, J. *et al.* In vivo organization of the FtsZ-ring by ZapA and ZapB revealed by quantitative super-resolution microscopy. *Mol Microbiol* **89**, 1099-1120, doi:10.1111/mmi.12331 (2013).

2 Bernhardt, T. G. & de Boer, P. A. The Escherichia coli amidase AmiC is a periplasmic septal ring component exported via the twin-arginine transport pathway. *Mol Microbiol* **48**, 1171-1182, doi:10.1046/j.1365-2958.2003.03511.x (2003).

3 Yang, X. *et al.* A two-track model for the spatiotemporal coordination of bacterial septal cell wall synthesis revealed by single-molecule imaging of FtsW. *Nat Microbiol* **6**, 584-593, doi:10.1038/s41564-020-00853-0 (2021).

4 Gonzalez, M. D., Akbay, E. A., Boyd, D. & Beckwith, J. Multiple interaction domains in FtsL, a protein component of the widely conserved bacterial FtsLBQ cell division complex. *J Bacteriol* **192**, 2757-2768, doi:10.1128/JB.01609-09 (2010).

5 Robichon, C., King, G. F., Goehring, N. W. & Beckwith, J. Artificial septal targeting of Bacillus subtilis cell division proteins in Escherichia coli: an interspecies approach to the study of protein-protein interactions in multiprotein complexes. *J Bacteriol* **190**, 6048-6059, doi:10.1128/JB.00462-08 (2008).

6 Wissel, M. C. & Weiss, D. S. Genetic analysis of the cell division protein FtsI (PBP3): amino acid substitutions that impair septal localization of FtsI and recruitment of FtsN. *J Bacteriol* **186**, 490-502, doi:10.1128/jb.186.2.490-502.2004 (2004).

7 Bryant, P., Pozzati, G. & Elofsson, A. Improved prediction of protein-protein interactions using AlphaFold2. *Nat Commun* **13**, 1265, doi:10.1038/s41467-022-28865-w (2022).

8 Morales Angeles, D., Macia-Valero, A., Bohorquez, L. C. & Scheffers, D. J. The PASTA domains of Bacillus subtilis PBP2B strengthen the interaction of PBP2B with DivIB. *Microbiology (Reading)* **166**, 826-836, doi:10.1099/mic.0.000957 (2020).

9 Bernardo-García, N. *et al.* Allostery, Recognition of Nascent Peptidoglycan, and Cross-linking of the Cell Wall by the Essential Penicillin-Binding Protein 2x of Streptococcus pneumoniae. *ACS Chemical Biology* **13**, 694-702, doi:10.1021/acschembio.7b00817 (2018).

10 Robichon, C., Karimova, G., Beckwith, J. & Ladant, D. Role of leucine zipper motifs in association of the Escherichia coli cell division proteins FtsL and FtsB. *J Bacteriol* **193**, 4988-4992, doi:10.1128/JB.00324-11 (2011).

11 Sjodt, M. *et al.* Structural coordination of polymerization and crosslinking by a SEDS-bPBP peptidoglycan synthase complex. *Nat Microbiol* **5**, 813-820, doi:10.1038/s41564-020-0687-z (2020).

12 Freischem, S. *et al.* Interaction Mode of the Novel Monobactam AIC499 Targeting Penicillin Binding Protein 3 of Gram-Negative Bacteria. *Biomolecules* **11** (2021).

13 Choi, Y. *et al.* Structural Insights into the FtsQ/FtsB/FtsL Complex, a Key Component of the Divisome. *Sci Rep* **8**, 18061, doi:10.1038/s41598-018-36001-2 (2018).

14 Brown, A. M. & Zondlo, N. J. A propensity scale for type II polyproline helices (PPII): aromatic amino acids in proline-rich sequences strongly disfavor PPII due to proline-aromatic interactions. *Biochemistry* **51**, 5041-5051, doi:10.1021/bi3002924 (2012).

15 Kashammer, L. *et al.* Divisome core complex in bacterial cell division revealed by cryo-EM. *bioRxiv*, 2022.2011.2021.517367, doi:10.1101/2022.11.21.517367 (2022).
